# Defining the transcriptional adaptation of *Staphylococcus aureus* to a range of nutritional sulfur supplementation

**DOI:** 10.64898/2026.05.20.726469

**Authors:** Paige J. Kies, Cristina Kraemer Zimpel, Joshua M. Lensmire, McKenna R. Major, Troy A. Burtchett, Michael R. Wischer, Neal D. Hammer

## Abstract

Bacterial pathogens must adapt to dynamic host tissue environments to proliferate. Accordingly, elegant regulatory systems evolved to overcome challenges presented by the host and satisfy nutritional requirements. Sulfur is an essential macronutrient and Gram-positive bacteria such as *Staphylococcus aureus* balance this nutritional requirement by employing the transcriptional repressor, CymR. Previous investigations defined the *S. aureus* CymR regulon by comparing transcripts generated in a *cymR* mutant cultured in cystine replete, rich medium to wild type cells. This study defines the *S. aureus* CymR-dependent and -independent sulfur-starvation response in chemically defined growth conditions. Results demonstrate that the sulfur starvation and sulfur replete CymR regulons exhibit considerable overlap, including previously noted connections between iron acquisition, oxidative stress, and sulfur metabolism. The link between iron acquisition, oxidative stress, and sulfur metabolism is validated further by the finding that sulfur-containing glutathione (GSH) mitigates heme and peroxide toxicity. In addition to GSH, Cys and thiosulfate fulfill the *S. aureus* sulfur requirement. Transcriptional responses to organic (cysteine, cystine, reduced and oxidized GSH) or inorganic thiosulfate were quantified, revealing sulfur source-specific expression patterns. Thiosulfate induced the largest number of differentially expressed genes. Consequently, the thiosulfate transporter (SAUSA300_RS10985) has been confirmed as essential for *S. aureus* growth when thiosulfate is the sulfur source. Furthermore, we demonstrate that a hypothetical protein operonic with SAUSA300_RS10985, SAUSA300_RS10980, supports maximal growth on thiosulfate. Collectively, a resourceful transcriptomics framework is provided which underscores the dynamic nature of *S. aureus* sulfur metabolism.

**IMPORTANCE:** The opportunistic pathogen *Staphylococcus aureus* proliferates within diverse host environments and consequently is a leading cause of hospital-acquired infections. During colonization *S. aureus* must acquire essential nutrients like sulfur to survive. Regulation of machinery to procure host-derived metabolites occurs to satisfy nutritional requirements and maintain homeostasis to circumvent detrimental impacts on bacterial physiology. This report investigates the *S. aureus* regulatory response during various states of sulfur supplementation *in vitro*, enhancing our knowledge of staphylococcal sulfur metabolism and represents a repository of information that will guide future research surrounding *S. aureus* sulfur acquisition and metabolism.

## INTRODUCTION

*Staphylococcus aureus* harbors considerable pathogenic potential given it adapts to and proliferates within most vertebrate organs (1, 2). Such capabilities demand bacteria sense and respond to the extracellular milieu to procure essential nutrients from the host. For example, regulatory mechanisms that control *S. aureus* transition metal acquisition and homeostasis are well defined (3). Transcriptional repressors, such as Fur, Zur, and MntR, modulate genes involved in acquisition and utilization of iron, zinc, and manganese, respectively. Each regulator has dedicated binding pockets for their cognate metal ion (i.e. Fe^2+^, Zn^2+^, and Mn^2+^) which senses intracellular concentrations, allowing the regulator to switch to an active, repressive conformation in replete environments (3). Activity of these metalloregulators is tightly coordinated given that transition metals, though essential, are detrimental when in excess (3–6).

Sulfur represents another essential nutrient whose coordinated acquisition by *S. aureus* involves similar regulation. Within cells sulfur is stored in the form of cysteine (Cys) which participates in numerous cellular reactions such as protein and co-factor synthesis or redox balance (6–13). Given this amino acid is the crux of sulfur distribution within a cell, all metabolites used to meet this nutritional requirement must be catabolized (organosulfur) or assimilated (inorganic sulfur) to Cys. As with iron, in *Escherichia coli* it has been demonstrated that an overabundance of intracellular Cys contributes to Fenton chemistry (14). Thus, stringent control of this element in coordination with iron is needed to accommodate proliferation (15). The major regulatory factor influencing *S. aureus* sulfur homeostasis is the <u>cy</u>steine <u>m</u>etabolism repressor, CymR (16–18). In contrast to the metalloregulators, CymR indirectly senses Cys levels to modulate DNA binding activity via two methods (Fig. 1). Intracellular Cys levels are communicated through the sulfur assimilation intermediate *O*-acetylserine (OAS). OAS quantities are inversely correlated with Cys. As Cys abundance increases, OAS is depleted, which causes CysK, an OAS-thiol-lyase or cysteine synthase, to respond by complexing with CymR, activating repression (Fig. 1A). Alternatively, CymR also responds to the oxidation status of the cell through its sole Cys residue at position 25 (Fig. 1B) (19). Recent work further demonstrated that CymR controls toxic levels of intracellular oxidized Cys, or cystine (CSSC), from impeding *S. aureus* proliferation in cystic fibrosis sputum (20). Decreased *cymR* mutant proliferation in the presence of abundant CSSC was alleviated when the CSSC transporters TcyP and TcyABC were genetically inactivated and the low molecular weight thiol antioxidant glutathione (GSH) was supplied as a sulfur source. These results highlight the critical role of CymR in maintaining growth-permissive levels of CSSC (20).

**Figure 1.**
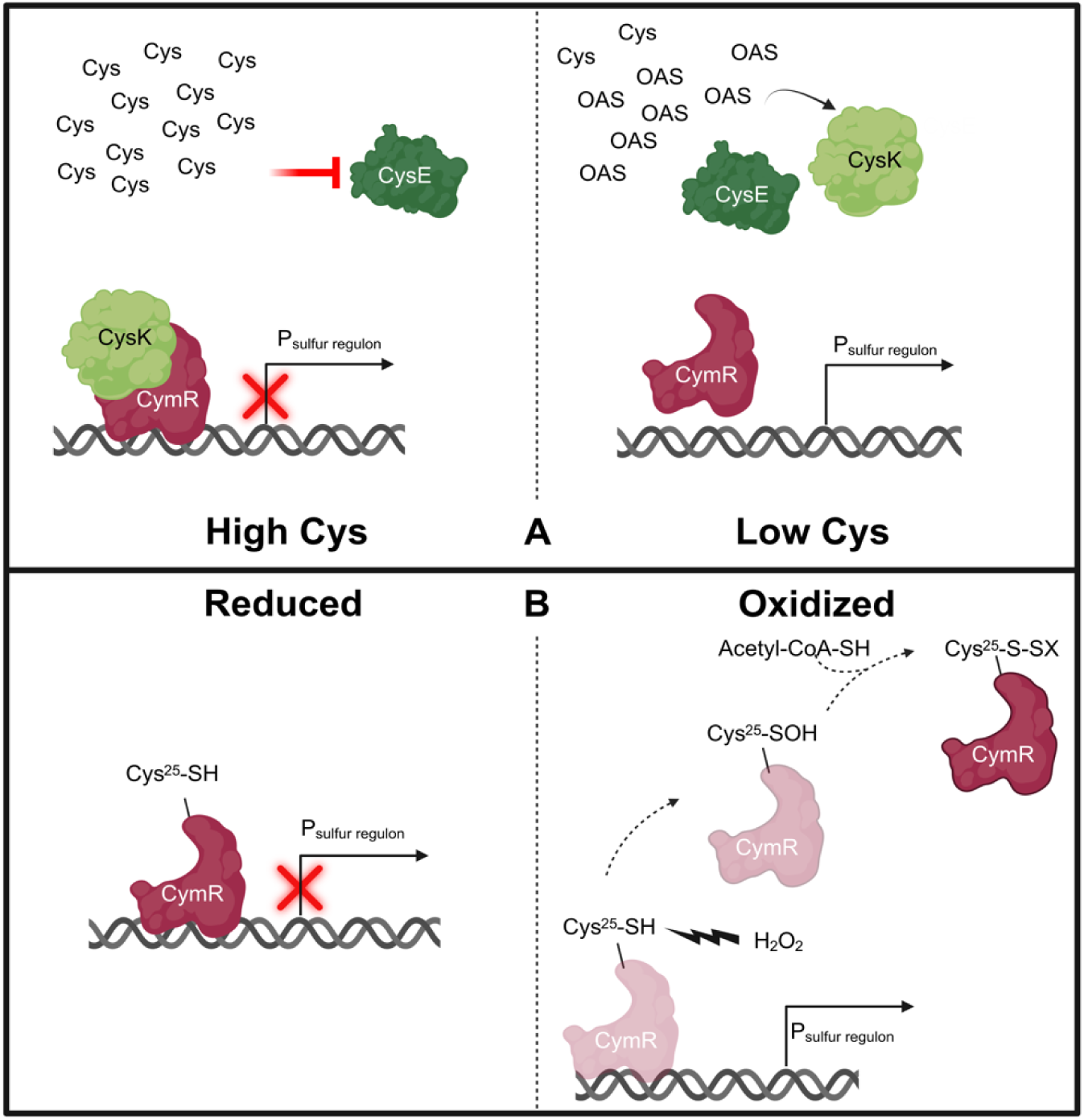
*S. aureus* CymR responds to at least two different stimuli. **(A)** Model of CymR function described by Soutourina *et al.* (18, 30, 31). CymR senses the intracellular Cys pool through the coordinated function of the serine transacetylase, CysE, the CysE product *O*-acetylserine (OAS), and cysteine synthase, CysK. CysE is inhibited by Cys when the amino acid is present at threshold concentrations (**A**, high Cys). In this situation, CysK interacts with CymR and the CysK-CymR complex binds target DNA sequences inhibiting expression of sulfur metabolism and acquisition genes. In the absence of Cys (**A**, low Cys), CysE produces OAS which is then consumed by CysK for use in inorganic sulfur assimilation. CymR repression is relieved and transcription of sulfur acquisition and metabolism genes proceeds. **(B)** An illustration of CymR sensing the oxidation state of the *S. aureus* cytoplasm via its sole Cys at position 25 (19). When Cys is in the reduced state (**B**, Reduced), CymR will bind to DNA. Upon Cys^25^ oxidation (**B**, Oxidized), the thiol group forms a sulfinic acid (-SOH) intermediate. The low molecular weight thiol, acetyl CoA then forms a disulfide bond with CymR, decreasing the affinity of the regulator for DNA. This illustration was created with BioRender.

In addition to Cys, CSSC, and GSH *S. aureus* can acquire oxidized GSH (GSSG) and thiosulfate (TS) to maintain intracellular sulfur and fulfill the elemental requirement (21–23). Together, these compounds participate in host sulfur metabolism across numerous tissues (16). Despite advances in our understanding of *S. aureus* sulfur metabolism, which includes characterization of the CymR regulon, how this organism responds to sulfur depletion and whether it prioritizes acquisition of distinct sulfur-containing metabolites remain outstanding questions.

This study sought to further define the *S. aureus* response to sulfur by establishing sulfur replete and deplete transcriptional profiles that emerge in a CymR-dependent and -independent manner. Using a chemically defined medium (CDM) supplemented with various sulfur sources, our findings expand upon work performed by Soutourina *et al*. which ascertained the *S. aureus* CymR regulon when cells were cultured in rich medium supplemented with 2 mM CSSC (18, 24). We observe a similar induction of iron transport and oxidative stress genes when *S. aureus* is sulfur starved (24). The importance of coupled regulation is established by demonstrating that *S. aureus* scavenging of exogenous nutrient sulfur in the form of GSH combats heme-induced oxidative stress as well as enhancing survival in response to hydrogen peroxide (H_2_O_2_), a naturally occurring biocide within vertebrate hosts. Comparing sulfur-source dependent transcriptional responses revealed that inorganic TS elicited the greatest number of differentially expressed genes compared to proliferation on the organic metabolites Cys, CSSC, GSH, and GSSG. Included in the differentially expressed genes we observed upregulation of SAUSA300_RS10985, a YeeE/YedE family transporter, and SAUSA300_RS10980 (25). We provide genetic evidence that these proteins are required for *S. aureus* proliferation on TS. The high degree of homology between SAUSA300_RS10985 and SAUSA300_RS10980 to the *E. coli* **t**hio**s**ulfate **u**ptake proteins <u>A</u> and <u>B</u> (TsuAB) supports the reannotating these genes *tsuA* and *tsuB*, respectively (25, 26). Overall, this work validates the importance of transcriptional links between sulfur metabolism, sulfur starvation, iron homeostasis, and oxidative stress responses as well as accentuating intricacies within the *S. aureus* sulfur regulon.

## RESULTS

### Integration of sulfur starvation and CymR transcriptomes refines the *Staphylococcus aureus* sulfur metabolism gene repertoire

A direct assessment of the *S. aureus* sulfur starvation response has not been performed. Previous work defined the CymR regulon using microarray hybridization of transcripts collected from a *cymR* deletion mutant (Δ*cymR*) generated in the methicillin susceptible SH1000 strain (MSSA) (18). The mutant and wild type (WT) strains were cultured in a rich medium (tryptic soy broth; TSB) supplemented with 2 mM cystine (CSSC). We sought to compare the sulfur starved transcriptome of WT *S. aureus* with a sulfur deprived *cymR* transposon mutant (*cymR*::Tn) to define the CymR-dependent and -independent response. As this assessment requires culture methods distinct from those employed previously, we first needed to establish conditions that induce *S. aureus* sulfur starvation. Soutourina *et al* reported that expression of the gene encoding the Cys and CSSC transporter, *tcyP*, exhibited the greatest increase in a *cymR* mutant (18, 22). In keeping with this, we generated a transcriptional reporter plasmid containing the *tcyP* promoter sequence upstream of YFP (pKK22-P*_tcyP_*-YFP). WT JE2, a laboratory derivative of the current endemic USA300 LAC strain, and an isogenic *cymR*::Tn mutant harboring pKK22-P*_tcyP_*-YFP were cultured in CDM supplemented with 25 µM CSSC to mid-exponential phase (27, 28). At this time, cells were washed and resuspended in fresh CDM supplemented with or without CSSC. An additional 2 h incubation was conducted to induce sulfur starvation. Fluorescence of *cymR*::Tn cells occurred regardless of the presence or absence of CSSC, confirming the *tcyP* promoter responds to CymR repression (Fig. 2). Compared to the sulfur replete CSSC-supplemented condition, WT cells displayed increased fluorescence after a 2 h incubation in sulfur depleted media (Fig. 2). These results reveal that *S. aureus* sulfur starvation is achieved after 2 h of sulfur depletion.

**Figure 2.**
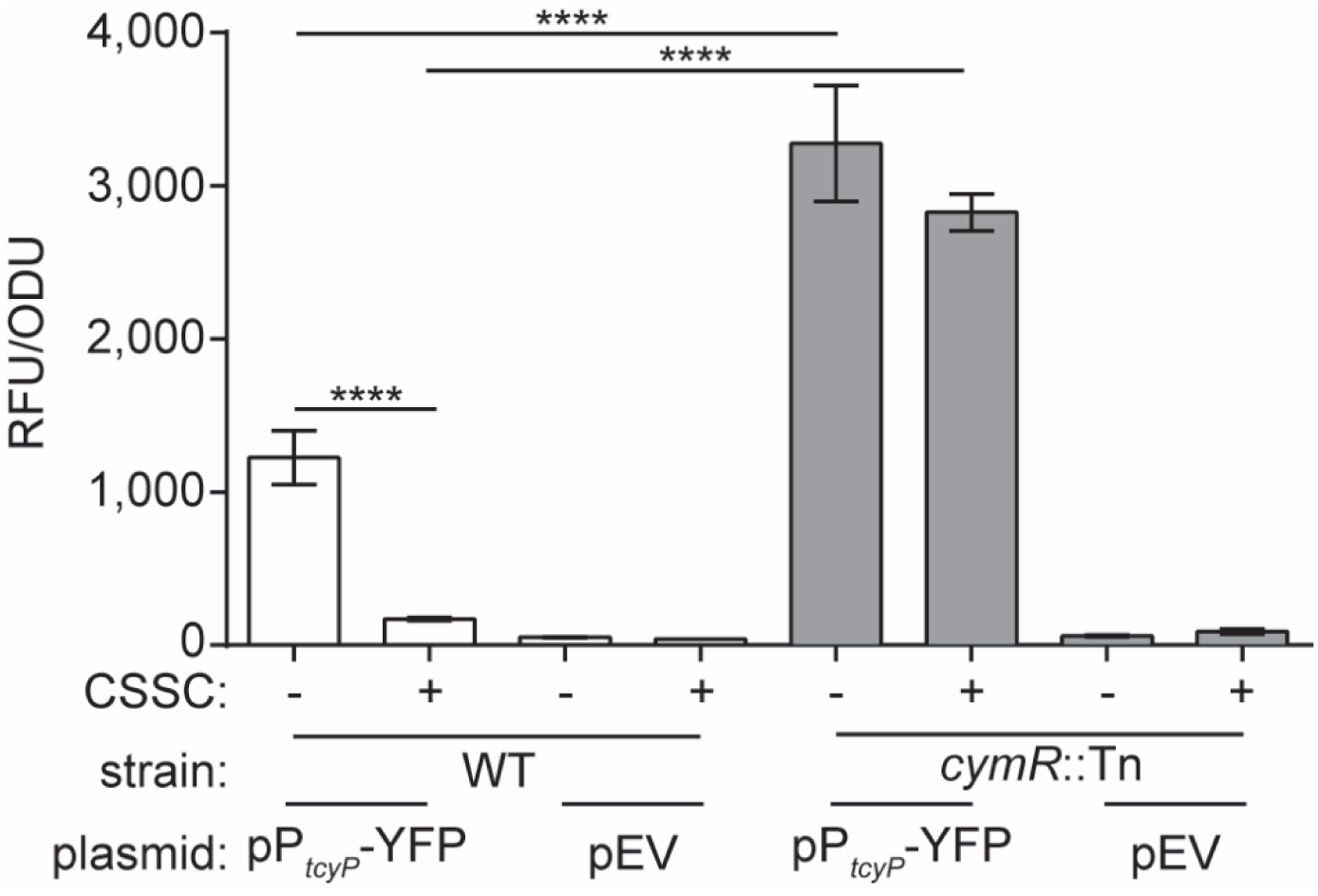
Quantifying induction of *S. aureus* sulfur starvation using a *tcyP* transcriptional reporter. WT (JE2) and isogenic *cymR*::Tn were cultured in chemically defined media (CDM) supplemented with 25 μM cystine (+CSSC) or lacking a sulfur source (-CSSC). Fluorescence and optical density from WT and *cymR*::Tn strains with or without the plasmid pKK22-P*_tcyP_*-YFP (pP*_tcyP_*-YFP) were measured. Strains harboring an empty vector (pEV) were used to monitor background fluorescence. The mean of three independent trials is presented and the error bars represent ±1 standard error of the mean. Statistical significance was calculated using the Welch ANOVA test. * Represents *P*-value <0.05, **** represents *P*-value <0.0001.

To determine the *S. aureus* response to sulfur starvation, RNAseq was performed using WT and the *cymR*::Tn mutant cells cultured in sulfur replete or deplete media (Fig. S1). Sulfur starvation was stimulated using culture conditions mimicking those that induced *tcyP* promoter activity (cells were cultured in CDM supplemented with CSSC to mid-log phase, washed and resuspended in CDM with or without CSSC and incubated for an additional 2 h to induce sulfur starvation; Fig. S1). Total numbers of differentially abundant transcripts across sulfur replete and sulfur deplete for WT and *cymR*::Tn (>2-fold, *P-*value < 0.05) are outlined in Table 1. In their initial study, Soutourina *et al*. identified 53 genes that altered expression at least 2-fold in a Δ*cymR* mutant (*P*-value ≥ 0.05, (18)). Included in their analysis were 16 genes associated with sulfur metabolism and acquisition. Using identical statistical cutoffs, a transcriptomic analysis of *cymR*::Tn cultured in sulfur replete CDM revealed that 62 genes changed abundance, 45 increased and 17 decreased (Table 1b, Dataset 2). Of the 16 CymR-regulated sulfur metabolism and acquisition genes reported by Soutourina *et al*., 10 were upregulated in the sulfur replete *cymR*::Tn mutant RNAseq dataset (Table 2, Fig.S2). Three of the six genes that missed the statistical cut-offs are operonic with genes that exhibited increased abundance (Fig. S2), including *tcyB* and *tcyC* that comprise the CSSC ABC transporter together with *tcyA* (22). Additionally, *mccA* (or *cysM*) and *mccB* (*metB*) reside within an operon and encode the enzymes that perform reverse transsulfuration of homocysteine to Cys (29–32). The remaining three genes that were previously identified but missed the statistical cut-off in this study include *ybdM* and two genes, SAUSA300_RS00935 and SAUSA300_RS00940, that are downstream of a putative sulfonate ABC importer (*ssuABC*, SAUSA300_RS00915 to SAUSA300_RS00925). Collectively, most but not all previously reported CymR-regulated genes were recovered, with missed targets likely reflecting statistical thresholding rather than biological exclusion. Importantly, based on the RNAseq results, the CymR regulon can be expanded further by incorporating genes that are functionally related and predicted to be co-transcribed with those upregulated in the *cymR::*Tn mutant. This includes the recently discovered GSH acquisition *gisABCD*-*ggt* operon and two putative ABC transporters, the one encoded by the aforementioned *ssuABC* operon and the other by SAUSA300_RS02330 to SAUSA300_RS02340 (apparent *metNPQ* homologues) (23). Taking these into account increases the number of *S. aureus* sulfur acquisition and metabolism genes regulated by CymR to 25 (Fig. S2). Of these 25 genes, 16 genes exhibited increased abundance in our sulfur replete *cymR*::Tn condition, including *tcyP* (Table 2, Table S1, and Dataset 2). Neither Soutourina *et al*. (16 out of 25) nor the sulfur replete *cymR*::Tn RNAseq dataset (16 out of 25) comprehensively capture the expanded 25-gene sulfur metabolism and acquisition network. Nonetheless, when combined with recent genetic experimentation (22, 23), the two data sets provide an expanded CymR regulated sulfur metabolism gene repertoire across strain backgrounds, RNA quantification approaches, and growth conditions.

**Table 1.** Number of differentially abundant transcripts associated with sulfur starvation, CymR regulation, and distinct sulfur sources.

| Culture comparison | Upregulated | Downregulated | Total DE genes<sup>a</sup> |
| --- | --- | --- | --- |
| WT -S vs WT +CSSC<sup>b</sup> | 425 | 121 | 546 |
| *cymR*::Tn +CSSC vs WT +CSSC<sup>c</sup> | 45 | 17 | 62 |
| *cymR*::Tn -S vs *cymR*::Tn +CSSC<sup>d</sup> | 170 | 80 | 250 |
| WT +Cys vs WT +CSSC | 4 | 2 | 6 |
| WT +GSSG vs WT +CSSC | 96 | 72 | 168 |
| WT +GSH vs WT +CSSC | 97 | 104 | 201 |
| WT +sTS vs WT +CSSC | 231 | 168 | 399 |

<sup>a</sup>Genes were differentially expressed (DE) with a Log<sub>2</sub> fold change equal to or greater than 1 with a *P*-value <0.05 from DESeq2 output
<sup>b</sup>Sulfur starvation response
<sup>c</sup>Sulfur-replete CymR regulon
<sup>d</sup>Starvation-induced changes independent of CymR

**Table 2.**
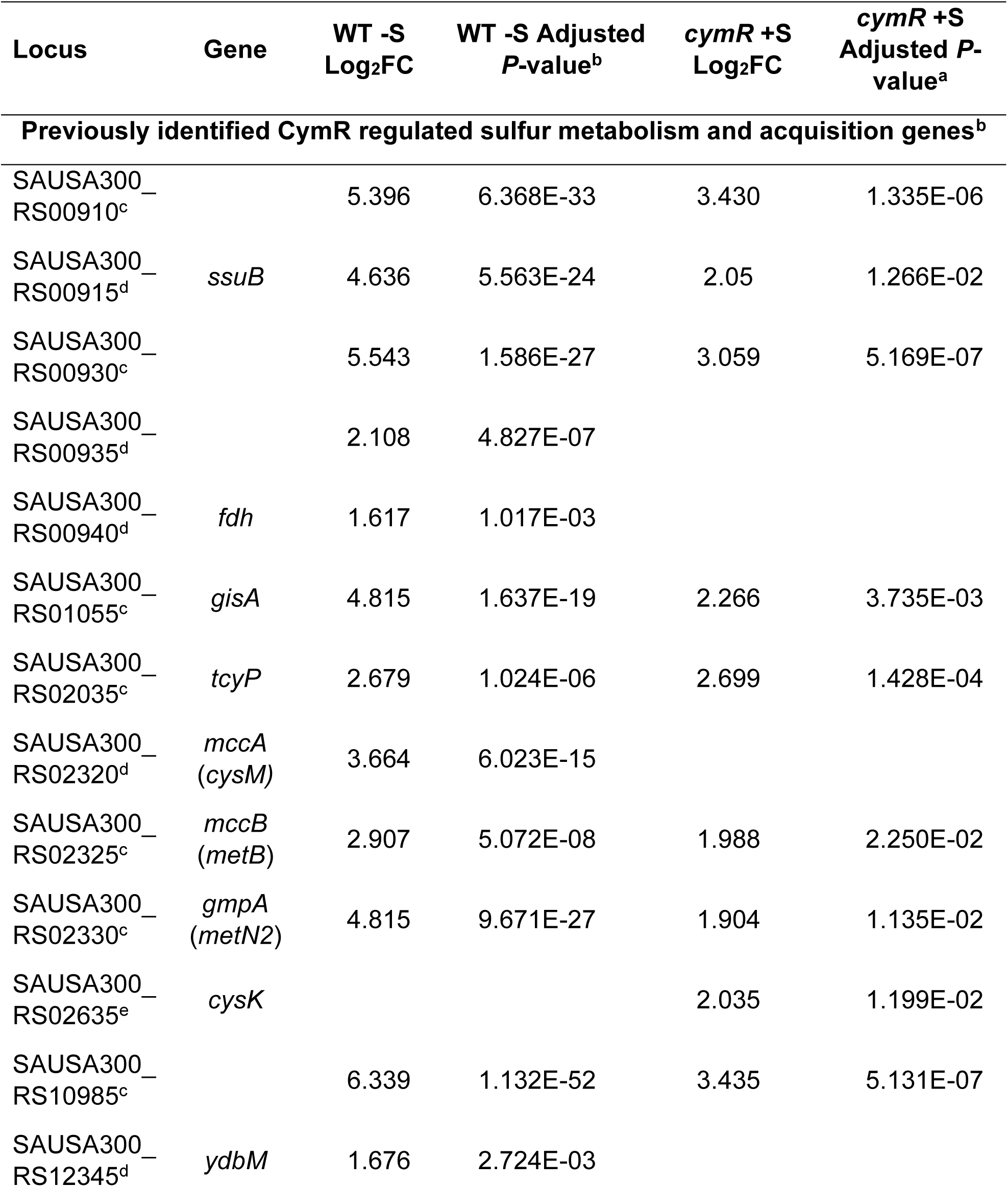
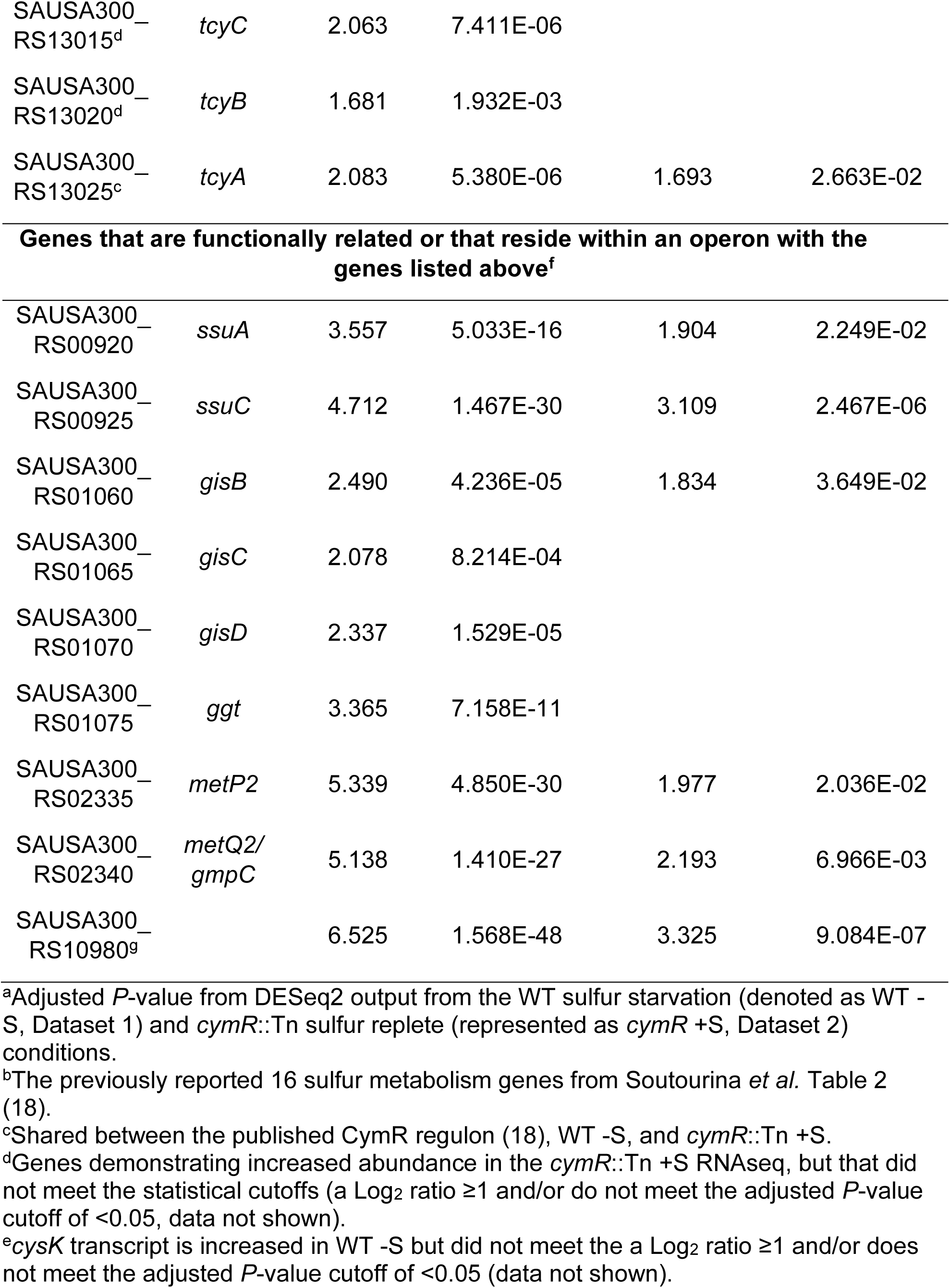
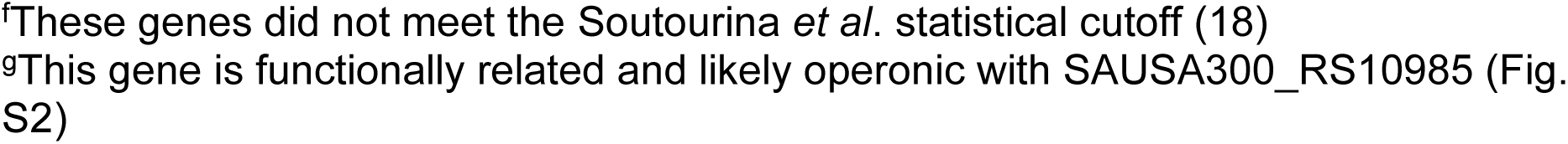
Comparative transcriptomics refines the *S. aureus* sulfur metabolism and acquisition gene repertoire.

CymR regulates gene expression based on internal Cys concentrations (Fig. 1) (32, 33), supporting the hypothesis that sulfur starvation activates the CymR regulon. Consistent with this, expression of 24 of the proposed 25 sulfur metabolism and acquisition genes are induced in the sulfur-starved WT condition. Only one of the previously recognized CymR-repressed genes, *cysK*, missed the statistical cutoff (Table 2, Fig. S2, and Dataset 1). Enrichment of CymR-regulated sulfur metabolism and acquisition transcripts in the WT sulfur starvation response demonstrate that the experimental approach and statistical analyses sufficiently capture the *S. aureus* sulfur starvation transcriptome. The findings also further reinforce the importance of CymR repression during sulfur deprivation. Overall, the WT sulfur starvation response included 425 transcripts that increased abundance and 121 that decreased abundance, resulting in a total of 546 differentially expressed transcripts (Table 1a and Dataset 1). The number of genes responding to sulfur starvation (*n* = 546) is greater than the previously described CymR regulon (*n* = 53, (18)) and the sulfur replete *cymR*::Tn transcriptome (*n* = 62). These results support the hypothesis that sulfur starvation induces CymR-dependent and -independent responses.

### The *S. aureus* sulfur starvation response is not exclusively governed by CymR

The disparity between the total number of genes differentially regulated in response to sulfur starvation (*n* = 546) and the sulfur replete CymR regulons (*n* = 53 and *n* = 62) underscores the importance of accounting for the sulfur status of cells. We surmise that CymR-dependent and -independent transcriptional alterations comprise the WT sulfur starvation response. Comparing WT and *cymR*::Tn provides a robust methodology to discern between the respective *S. aureus* CymR-dependent and - independent responses to sulfur starvation. Differentially abundant transcripts shared between sulfur-starved WT and *cymR*::Tn cells reflect CymR-independent regulation, as their expression changes irrespective of CymR status. Induction of sulfur starvation in *cymR*::Tn cells alters the abundance of 250 genes, 170 increase and 80 decrease (Table 1d and Dataset 3). Of those 250 genes, 133 are also differentially regulated in sulfur starved WT cells, indicating a substantial portion of the sulfur starvation response is regulated independently of CymR (Fig. S3A, comparison between Datasets 1 and 3). Genes of unknown function represent the largest COG category within the sulfur starved CymR-independent regulon (Fig. S3B, Fisher’s test for enrichment *P* = 0.256).

While these 133 genes represent the CymR-independent transcriptional response to sulfur starvation the remaining 413 genes are CymR-dependent, highlighting the dominance of CymR in driving the sulfur starvation response. COG analysis revealed early 200 genes in the WT sulfur starved condition were assigned to either unknown function (Fisher’s test for enrichment *P* < 0.001) or no homologue found (Fisher’s test *P* = 0.086) indicating there is much to be learned about the *S. aureus* response to sulfur starvation (Fig. S3B). Lastly, a subset of genes exhibited differential abundance solely in the *cymR*::Tn sulfur starvation response (*n* = 117, Fig. S3A). Though these genes are considered CymR-independent, their differential expression is likely a compensatory response to the loss of CymR. The most prevalent COG categories within this subset include genes of unknown function (Fisher’s test *P* = 1.0) and amino acid transport and metabolism (Fisher’s test *P* = 0.021, Fig. S3B). Together, these comparisons uncover a multifaceted *S. aureus* sulfur starvation response that is largely CymR-dependent but also includes CymR-independent transcriptional alterations within and beyond the sulfur metabolism repertoire.

### Iron acquisition and oxidative stress response genes are induced during *S. aureus* sulfur limitation

Subsequent to their initial CymR paper, Soutourina *et al*. published a second study that demonstrated increase abundance of 18 oxidative stress and metal ion homeostasis genes when the Δ*cymR* mutant was grown in TSB supplemented with 2 mM CSSC (18, 24). Our results further corroborate a link between CymR, iron acquisition, oxidative stress and sulfur deprivation. Of the 546 differentially expressed genes in sulfur starved WT (Dataset 1), 33 encode transcriptional regulators, of which 28 increased abundance and 5 decreased (Table S2). Notably, genes encoding the hallmark transcriptional regulators Fur and PerR that coordinate the iron acquisition and the peroxide stress responses, respectively, exhibited increased abundance (Table S2). Activation of Fur and PerR is further supported by the number of genes within the respective regulons that increased abundance. Of the 45 Fur-regulated genes, 29 exhibited increased abundance upon sulfur starvation, and abundance of six of the eight PerR-regulated genes also increased (the Fisher’s exact test *P* values for both the Fur and PerR regulons is < 0.001) (34–37). Increased *fur* and *perR* transcript abundance were also observed in Soutourina *et al*. (24). Interestingly, increased *fur* transcript abundance was observed in both sulfur-starved WT and the *cymR*::Tn mutant (Dataset 1 and Dataset 3), despite the inclusion of iron in the defined medium. The fact that iron acquisition genes increase abundance in both the WT and *cymR*::Tn sulfur starvation conditions indicate that their induction is CymR-independent (Table 3, asterisk). However, several iron acquisition genes are uniquely abundant in the WT condition. For example, genes encoding the siderophore importer FhuB and FhuG increase abundance in WT but not *cymR*::Tn, suggesting their expression is controlled by CymR (Table 3) (38). On the other hand, 14 genes involved in Fur-regulated siderophore biosynthesis and Fe-S cluster assembly, are unique to the *cymR*::Tn starvation condition. Included in this group is the regulator of redox-stress HypR (SAUSA300_RS03085; Table 3) (39). Other transcriptional regulators exhibiting differential abundance in the sulfur starved *cymR*::Tn mutant include the virulence regulator *sarR* (SAUSA300_RS12390) (40, 41) which is downregulated, while SAUSA300_RS11905, a putative MerR family of metal sensing regulators displayed increased abundance (40, 42). These CymR compensatory transcriptional changes highlight the activation of alternative regulatory pathways associated with iron acquisition, oxidative stress, and metal ion homeostasis under sulfur-limiting conditions.

**Table 3.**
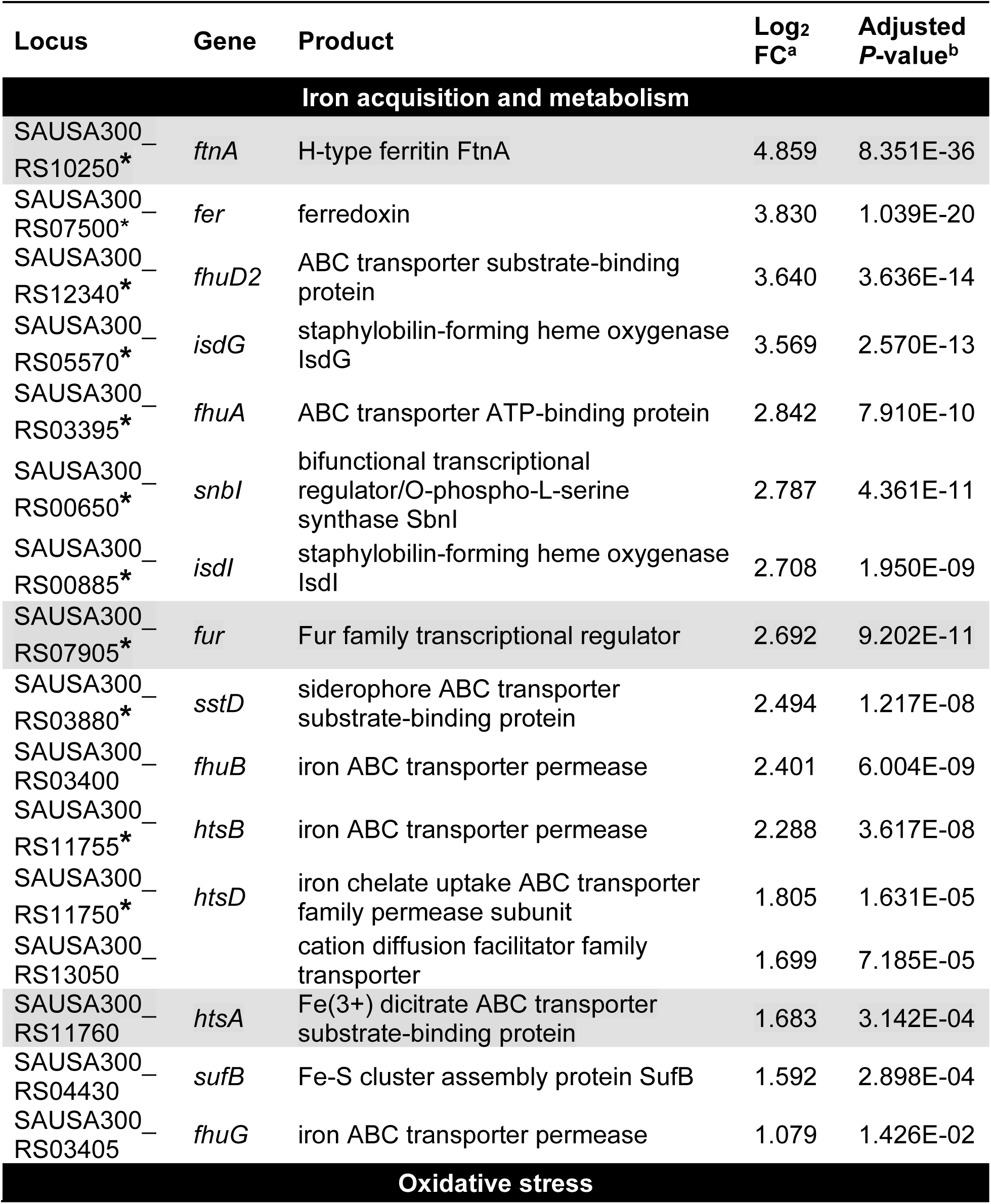
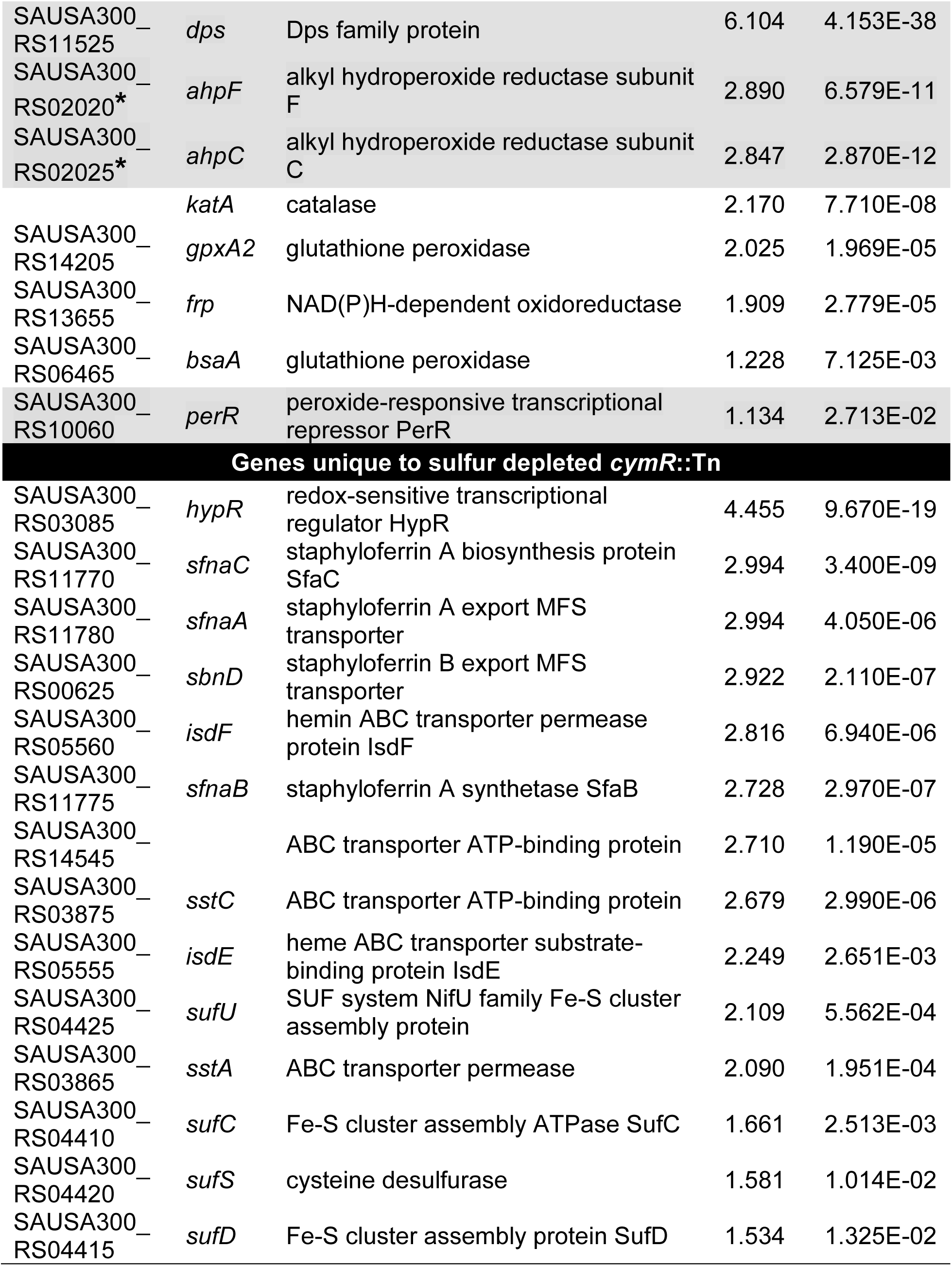
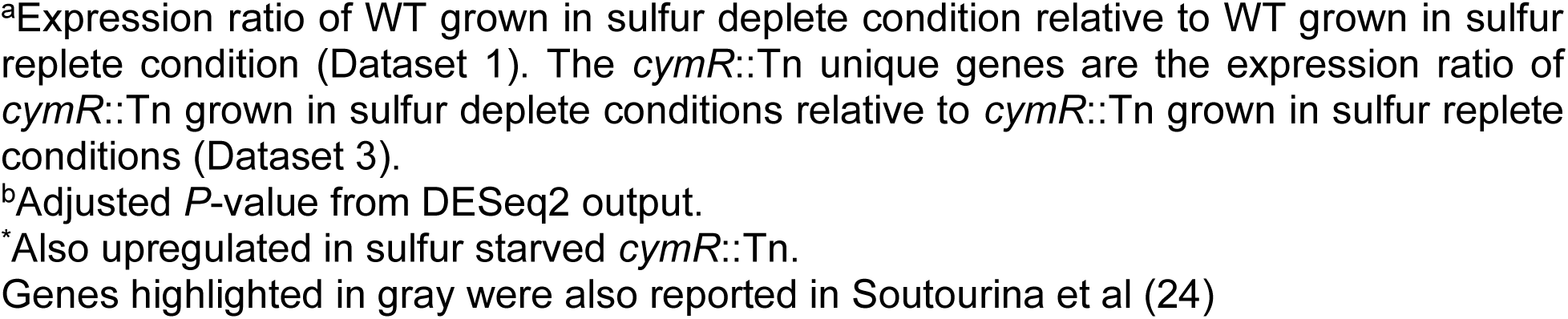
*S. aureus* sulfur starvation induces expression of iron acquisition and oxidative stress response genes.

The observed interconnection between sulfur, iron, and oxidative stress likely benefits *S. aureus* because numerous enzymes require Fe-S clusters to function and homeostasis of these two nutrients is crucial given their contributions to Fenton chemistry which damages Fe-S clusters (14, 43, 44). Previous work has shown that exogenous GSH protects *Streptococcus* species against oxidative damage upon exposure to copper or paraquat (45, 46). Together, these facts support the hypothesis that *S. aureus* scavenges host antioxidant thiols, like GSH, to not only meet nutritional demands, but to also manage oxidative stress. Given the Isd system is important for heme-mediated iron acquisition and the fact that erythrocytes contain approximately 1-2 mM GSH and 20 mM heme (20 µM of which is predicted to be dynamically ‘free’ heme, (44, 47–50)), we explored *S. aureus* sensitivity to hemin (oxidized heme) upon supplementation with reduced or oxidized GSH. WT and a *hrtA*::Tn mutant, which is hypersensitive to heme due to impaired efflux (51), displayed identical growth profiles when cultured in CDM supplemented with 50 μM GSH (Fig. 3A). However, toxicity is observed as an increased lag phase upon supplementation of the WT culture with 10 μM hemin, while the *hrtA*::Tn mutant fails to proliferate (Fig. 3A). To reflect GSH levels *S. aureus* likely encounters during infection, GSH was increased to a more physiologically relevant concentration (49, 50); supplementation with 750 μM GSH abolished any impact of hemin on WT propagation (Fig. 3B). The *hrtA* deficient strain experienced a longer lag phase but eventually reached a WT-like OD_600_ (Fig. 3B), indicating an alleviation of heme toxicity. To assess whether this protection is dependent on free thiol, reduced GSH was replaced with oxidized GSH (GSSG) using equimolar concentrations of sulfur. Exposing S. *aureus* supplemented with low (25 μM) or high (375 μM) concentrations of GSSG to hemin inhibited growth of both WT and the *hrtA* deficient strain (Fig. 3C, D respectively). Taken together, this indicates that *S*. *aureus* scavenging of GSH sufficiently combats hemin-mediate oxidative stress. To further demonstrate that GSH protects against ROS, we quantified the viability of *S. aureus* cells that were preincubated with GSH or GSSG and subsequently exposed to H_2_O_2_ (Fig. 3E). WT cultured in 750 μM GSH exhibited significantly increased viability upon H_2_O_2_ challenge compared to WT grown in 50 μM GSH or 375 μM GSSG. Collectively, these data indicate that sulfur starvation and iron homeostasis coordination protect *S. aureus* from oxidative stress.

**Figure 3.**
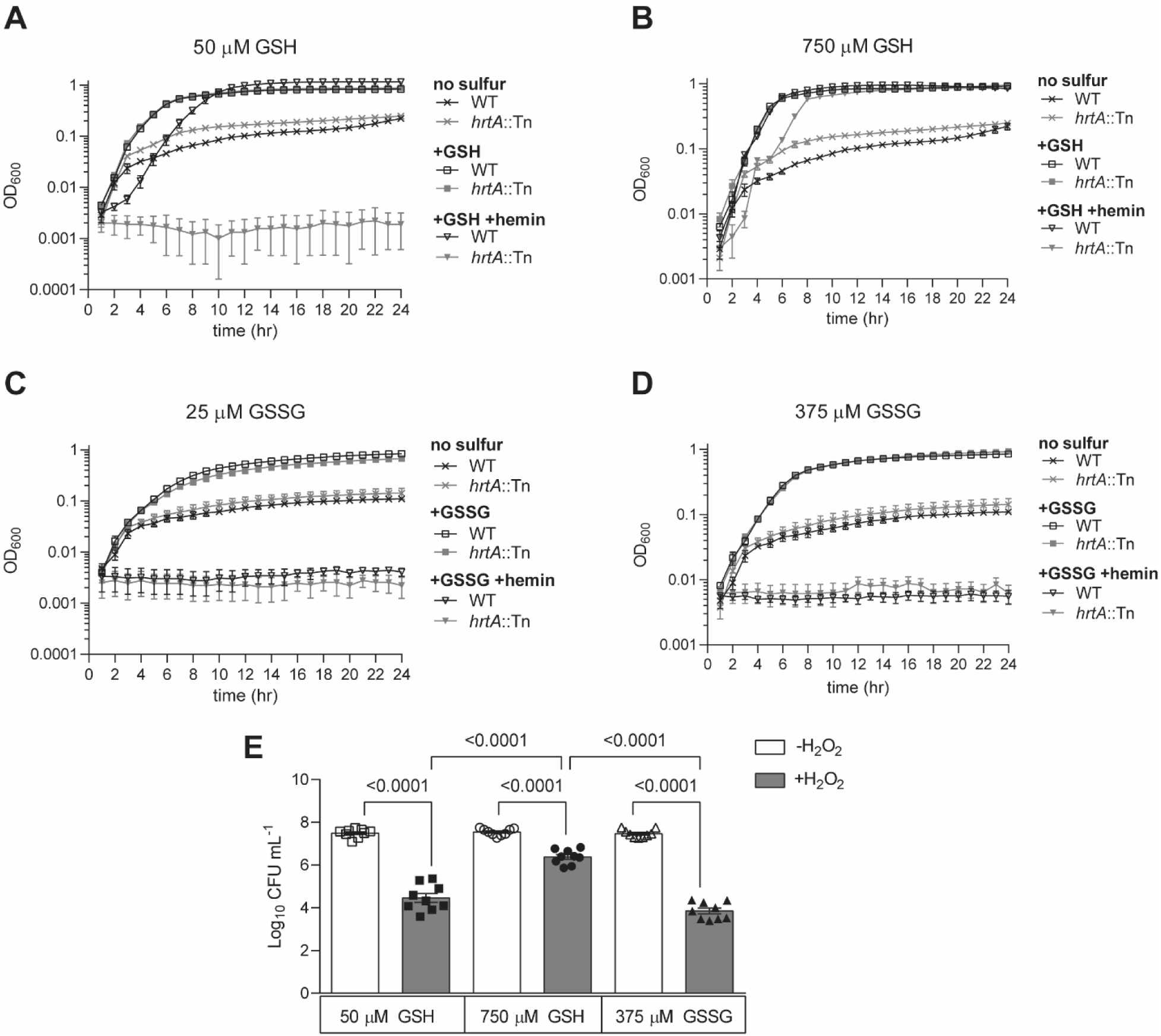
GSH alleviates oxidative stress in *S. aureus*. WT and *hrtA*::Tn were cultured in chemically defined medium (CDM) in the absence of a sulfur source or supplemented with 50 µM or 750 µM reduced glutathione (GSH) (**A** and **B**, respectively) as well as 25 µM or 375 µM oxidized glutathione (GSSG) (**C** and **D**, respectively) in the presence or absence of 10 μM hemin. Curves are the mean of three independent trials and the error bars represent ±1 standard error of the mean. **(E)** WT preloaded with either 50 μM GSH (squares), 750 μM GSH (circles), or 375 μM GSSG (triangles) were left untreated (open symbols) or exposed to 1 M H_2_O_2_ (closed symbols) prior to CFU enumeration. Presented are the average of three independent trials ±1 standard error of the mean. Statistical significance represents Brown-Forsythe and Welch ANOVA tests. * Represents *P*-values <0.05 while ** represents *P*-values <0.005. ns denotes not significant indicating a *P*-value >0.05.

### Adaptation of *S. aureus* to distinct nutrient sulfur sources induces unique transcriptional responses

We next sought to define the transcriptional response of *S. aureus* to distinct sulfur sources by culturing the bacteria in CDM supplemented with 50 µM Cys, 25 µM GSSG, 50 µM GSH, or 50 µM sodium thiosulfate (sTS) (Datasets 4, 5, 6, and 7, respectively). CSSC was used as the comparator because it is typically supplemented as the sulfur source in the defined medium (27, 28). In total, 774 genes alter abundance when *S. aureus* proliferates in the presence of Cys, GSSG, GSH, or sTS compared to CSSC (Datasets 4-7). Only six genes were differentially abundant in WT supplemented with Cys (Table 1, Dataset 4). One of the two genes displaying decreased abundance in the Cys-supplemented condition, SAUSA300_RS04580, encodes a putative pyridine nucleotide disulfide oxidoreductase (40), suggesting a role in reducing CSSC. However, inactivation of SAUSA300_RS04580 does not impact the ability of *S. aureus* to utilize CSSC as a sulfur source (Fig. S4).

In stark contrast to CSSC, sTS supplementation led to the greatest transcriptional shift as 231 genes increased abundance while 168 decreased abundance (Table 1, Fig. 4A). Compared to growth in GSH and GSSG, sTS supplementation resulted in 135 genes increased and 76 genes decreased in abundance (Fig. 4B and C). Supplementation with GSSG altered the abundance of 168 genes (96 increased and 72 decreased; Table 1, Dataset 5), while GSH supplementation altered the abundance of 201 genes (97 increased and 104 decreased, Table 1, Dataset 6; Fig. 4A). A possible explanation for the vast transcriptional fluctuations in response to distinct sulfur sources is the differential abundance of transcripts that encode transcriptional regulators. In total, transcript abundance of 18 regulators changed. Of these, seven have been experimentally validated (36, 52–57). A majority change in response to sTS (n = 17), five upon GSH supplementation, and three in response to GSSG. Eleven of these regulators were solely altered in sTS (Fig. 4D). Only one regulator, *vraR* (SAUSA300_RS10185), displayed decreased transcript abundance across all three conditions (Fig. 4D). The 11 regulators uniquely expressed in sTS support the notion that altered gene expression in the presence of this sulfur source is related to transcriptional regulation. Overall, these observations indicate that growth on inorganic sTS alters the *S. aureus* transcriptional profile to a greater extent than when proliferation is supported by organic sulfur sources.

**Figure 4.**
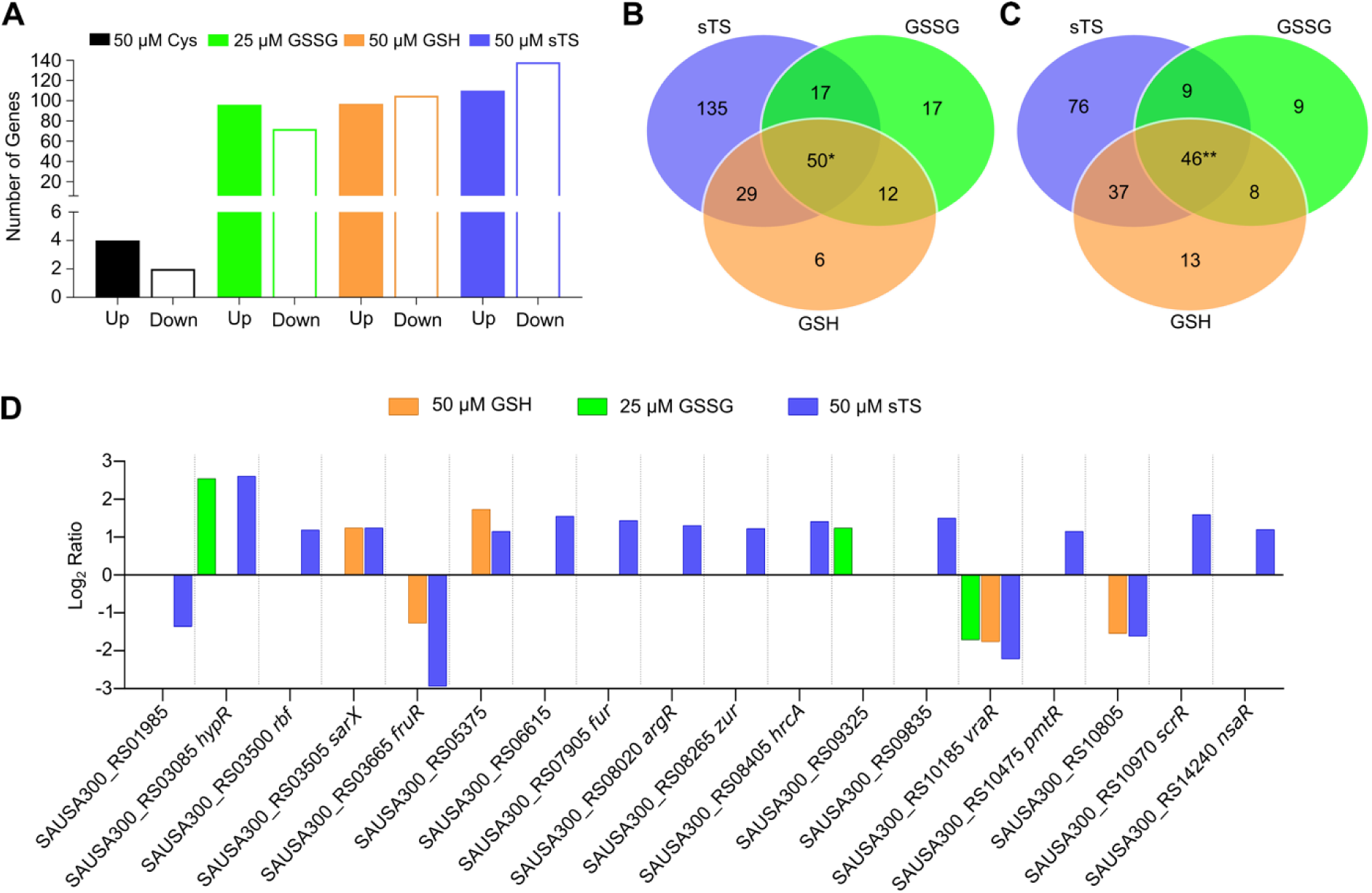
Supplementation with sTS induces considerable transcriptional changes in *S. aureus*. Differentially abundant transcripts in WT cells were identified by comparing cells cultured in chemically defined medium (CDM) supplemented with cysteine (Cys; black), oxidized glutathione (GSSG; green), reduced glutathione (GSH; orange), or sodium thiosulfate (sTS; purple) to CDM supplemented with cystine (CSSC). **(A)** Total number of genes that are differentially expressed in response to distinct sources of nutrient sulfur. **(B)** Venn diagram comparing genes upregulated when *S. aureus* is cultured in GSSG-, GSH- or sTS-supplemented medium. *The four genes exhibiting increased abundance in CSSC (SAUSA300_RS15735, SAUSA300_RS15740, SAUSA300_15090, and SAUSA300_RS15730) were not shared with GSSG, GSH, or sTS and were thus excluded from the Venn Diagram. **(C)** Genes displaying decreased abundance between the GSSG, GSH, and sTS. *Two genes decreasing in abundance in CSSC (SAUSA300_04580 and SAUSA300_06690) were both shared with GSSG, GSH, and sTS and are represented within the total shared genes in the Venn Diagram. **(D)** Transcriptional regulators differentially abundant when *S. aureus* is grown on GSH, GSSG or sTS compared to CSSC.

The putative YedE/YeeE family transporter SAUSA300_RS10985 (referred to as 985 hereafter) was captured across all sulfur source supplementation datasets and exhibited increased abundance in the WT sulfur starvation transcriptome as well as the CymR regulons (Table 2, Fig. S2, (18)); however, the substrate for this transporter in *S. aureus* is not known. An *E. coli* homologue of 985, TsuA, was found to import sTS (25). TsuA is operonic with TsuB a putative oxidoreductase that also supports *E. coli* TS utilization (25, 26). Similarly, the putative TsuB homologue in *S. aureus*, SAUSA300_RS10980 (referred to as 980 hereafter) is predicted to be co-transcribed with 985. The homology, gene expression pattern, and gene orientation support the hypothesis that 980 and 985 facilitate *S. aureus* TS utilization as a sulfur source (19, 58). To test this, a 985::Tn mutant was generated and assessed. The Tn insertion likely has polar effects on the downstream 980 gene (Fig. 5A). As such, complementation was performed using the pKK22 vector (59). Both genes were cloned, under control of the promoter, into pKK22 individually or as an operon. The resulting strains were then cultured in CDM supplemented with either 50 µM sTS or 25 µM CSSC (Fig. 5B, C respectively). In comparison to WT harboring an empty vector (EV), the 985::Tn EV mutant failed to proliferate when sTS is the sulfur source (Fig. 5B). Expression of 985 *in trans* partially restored propagation of the mutant while complementation with the entire operon rescued the observed growth defect. Ectopic expression of 980 did not restore proliferation of the mutant in CDM supplemented with sTS. These results demonstrate that 985, a putative transporter, is necessary for maximal growth of *S. aureus* on sTS while 980 performs an auxiliary role. Each phenotype is specific to sTS given that the mutant strains demonstrate WT-like growth on the control sulfur source, CSSC (Fig. 5C). In accordance with the nomenclature proposed by Morigasaki *et. al*. we propose to designate 985 and 980 thio<u>s</u>ulfate <u>u</u>ptake proteins <u>A</u> and <u>B,</u> TsuA and TsuB, respectively (25, 60).

**Figure 5.**
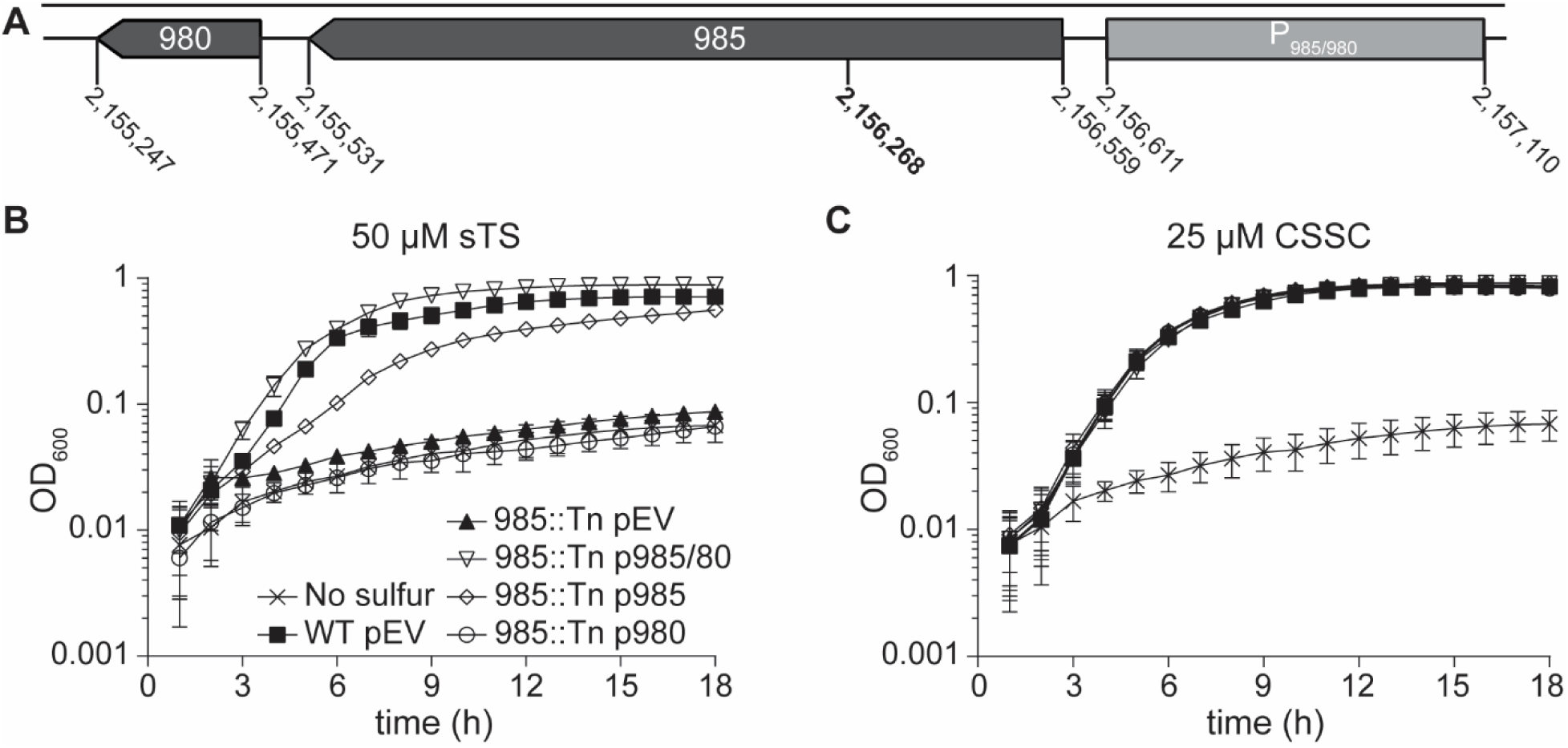
*S. aureus* employs TsuAB to utilize thiosulfate as a source of nutrient sulfur. **(A)** An illustration of the 985 (SAUSA300_RS10985) and 980 (SAUSA300_RS10980) operon and location of the Tn insertion (position 2,156,268, bold) (60). P_985/80_ (position 2,156,611-2,157,110) denotes the locus of the native promoter used in complementation studies. **(B-C)** Resulting growth kinetics of strains cultured in chemically defined medium (CDM) supplemented with either 50 µM sodium thiosulfate (sTS) **(B)** or 25 µM cystine (CSSC) **(C)** as the sole sulfur source. WT *S. aureus* harboring an empty pKK22 vector (pEV) cultured in medium lacking a viable sulfur source represents the no sulfur control. The mean of three independent trials is presented and the error bars represent ±1 standard error of the mean.

## DISCUSSION

As knowledge regarding *S. aureus* sulfur acquisition and assimilation mechanisms expands, it is vital to understand how these strategies are regulated (22, 23). Previous publications defined the *S. aureus* CymR regulon using an isogenic mutant cultured in sulfur replete media (18). Here, we examine the CymR-dependent and CymR-independent transcriptomes under sulfur replete and deplete conditions using RNAseq. WT and *cymR*::Tn cells were cultured in a defined medium in the presence or absence of CSSC allowing us to determine the sulfur starvation transcriptome. Our study also assessed a methicillin resistant, USA300 LAC derivative (JE2) which is reflective of the current endemic strain, whereas previous work utilized the methicillin susceptible, laboratory derived strain SH1000. Comparing our WT sulfur starvation to the sulfur replete CymR transcriptomes using identical statistical cutoffs, *P*-value of <0.05 using both Wilcoxon and Z-tests that showed a ≥2-fold change in transcript levels (18), revealed a greater number of differentially abundant genes (Table 1). This can be attributed to the fact that CymR-dependent and -independent responses contribute to the *S. aureus* sulfur starvation adaptation. Several other factors likely contribute to disparities between the Datasets presented here and the previously reported CymR regulon including increased sensitivity of RNAseq compared to microarray hybridization, culturing cells in CDM compared to rich media, and variations between strain backgrounds in methicillin resistant and methicillin susceptible *S. aureus* strains.

Despite differences in the total number of differentially abundant genes, similar induction of genes associated with sulfur metabolism and transporters was observed between the CymR regulons and WT sulfur starved Datasets (sulfur-starved WT Dataset 1; CSSC-supplemented *cymR*::Tn mutant Dataset 2; Table S1; and the CymR regulon described in Soutourina *et al*.) (18). For example, increased transcript abundance of one of the two Cys and CSSC transporters, TcyP (SAUSA300_RS02035) was consistent across all three Datasets and Soutourina *et al* (18, 22). Similarly, transcripts corresponding to the lipoprotein component of the Cys/CSSC ABC transporter TcyABC (SAUSA300_RS13015-RS13025), *tcyA* was also increased across the three conditions. However, the remaining *tcyB*-encoded permease and ATPase encoded by *tcyC* were only upregulated in WT sulfur starvation and Soutourina *et al*., but not in the CSSC replete *cymR*::Tn mutant. Heightened expression of a recently described GSH import system (*gisABCD-ggt,* SAUSA300_RS01055-RS01075) was also observed across the three analyses (23). The ABC transporter is encoded by divergently transcribed *gisA* and *gisBCD*. The terminal gene of the *gisBCD* operon is *ggt*, which encodes a γ-glutamyl transpeptidase. All five genes exhibit increased abundance in sulfur starved WT. Both *gisA* and *gisB* are expressed in CSSC supplemented *cymR*::Tn while Soutourina *et al*. observe upregulation of only *gisA* (Fig. S2) (18). The genes encoding for a predicted sulfate/sulfonate transport system, *ssuABC* (SAUSA300_RS00915 to SAUSA300_RS00925), were also more abundant in both the CSSC supplemented *cymR*::Tn and WT sulfur starvation conditions; however, only *ssuB* was upregulated in Soutourina *et al*. Despite these subtle inconsistencies across operonic genes, expression of the *gisABCD-ggt* and *ssuABC* loci further solidifies the overlap within the compared Datasets. Lastly, increased abundance of *tsuA*, the verified TS transporter, occurred across all three Datasets. Comparing these transcriptional profiles provides a more robust interpretation of CymR-dependent and sulfur-responsive networks governing sulfur metabolism and acquisition in *S. aureus*. This analysis culminates in a revised set of 25 genes that contribute to *S. aureus* sulfur metabolism and acquisition (Table 2, Fig. S2).

A portion of the sulfur starvation response involves genes outside of the CymR regulon (Fig. S3) such as those involved in iron metabolism and oxidative stress (Table 3 and Dataset 4). A potential link between iron and sulfur regulons seems intuitive considering the number of enzymes that require Fe-S clusters (14, 43, 44). In *Pseudomonas aeruginosa* the sulfur regulon transcriptional regulator, CysB, directly promotes expression of *pvdS*, an alternative sigma factor involved in the iron response (61), suggesting beneficial effects of balancing iron and sulfur levels. This coordination is further exemplified in *E. coli* Fur which has the capacity to respond directly to [2Fe-2S] clusters that activate repression (15); whether Fur is also responsive to CysB in this organism, though, is currently unknown. The fact that iron and sulfur potentiate Fenton chemistry further accentuates coordination between sulfur acquisition, iron acquisition, and oxidative stress.

Bacterial scavenging of host GSH, a low molecular weight thiol antioxidant, to satisfy the nutritional sulfur requirement potentially offsets toxicity associated with iron or heme-iron acquisition during infection. We demonstrate that GSH protects *S. aureus* from hemin toxicity, supporting propagation in an otherwise inhibitory environment (Fig. 3). To combat noxious heme accumulation, *S. aureus* employs the heme-regulated ABC transporter, HrtAB, which effluxes heme from the cell. Therefore, genetic inactivation of *hrtA* or *hrtB* results in heme hypersensitivity (51, 62). Here we observe that GSH scavenging promotes proliferation of both WT and a *hrtA*::Tn mutant in the presence of hemin. The free thiol of GSH is a crucial aspect of this protection given that GSSG is not sufficient to alleviate the toxicity. Additionally, we demonstrate that H_2_O_2_ toxicity is diminished upon culturing *S. aureus* in the presence of physiologically relevant GSH concentrations. This supports the notion that the restored growth observed in response to heme is due to management of cell-associated stress induced by heme rather than an extracellular GSH-heme interaction (63, 64). The relevance of these results towards *S. aureus* management of heme-induced stress during bacteremia stem from the facts that the pathogen is hemolytic, and erythrocytes harbor abundant concentrations of GSH and heme (65). Thus, coordination of the sulfur, iron, and oxidative stress regulons benefit *S. aureus*.

Due to the numerous metabolites that satisfy the *S. aureus* sulfur requirement, it is unknown whether this pathogen encounters sulfur-limited environments during infection. *S. aureus* has evolved several elegant mechanisms to pillage host-derived sulfur reservoirs (16). Sulfur-containing metabolites vary in abundance or composition depending on tissue types and sub-cellular localization (16). Given this, it is accordingly plausible that *S. aureus* adapts to the local sulfur milieu by altering its sulfur acquisition and metabolism transcriptome. Growth of *S. aureus* on four different sulfur sources (Cys, GSH, GSSG, and sTS) reflects this. In comparison to the CSSC condition, each sulfur source resulted in at least 6 (Cys) and a maximum of 399 (sTS) differentially abundant transcripts.

A limitation of the current study and previous published CymR regulon is single time point assessment of the transcriptome. We observed induction of the sulfur starvation regulon upon supplementation with GSH, GSSG, and sTS. Despite the presence of an adequate sulfur source, *S. aureus* was primed to import other sulfur-containing metabolites. Future studies focused on quantifying kinetic expression of the transporters while simultaneously monitoring OAS levels and CymR ChIPseq will provide a better understanding for why sulfur transporters are expressed despite supplementation with adequate nutrient sulfur. Finally, the Δ*cymR* mutant used in the Soutourina *et al*. study exhibited decreased hemolytic activity and virulence though differential expression of virulence factors was not reported (18, 33). Similarly, our conditions did not reveal an overabundance of differentially expressed virulence genes. We attribute this to harvesting RNA from mid-exponential cells when Agr, a major virulence regulator, activity is low.

Collectively, our investigation into the *S. aureu*s sulfur response reveals intricacy surrounding CymR and the sulfur regulon. This manifests when examining the molecular mechanisms of TS assimilation. Extensive characterization of TS assimilation in *E. coli* and *Salmonella typhimurium* has generated a model where the CysPUWA ABC transporter is the primary TS importer (63, 66, 67). *S. aureus* does not encode a CysPUWA homolog and recent work in *E. coli* identified the TsuAB (40). Here we demonstrate that *S. aureus* employs TsuA to grow on TS (Fig. 5B). Additionally, TsuB is thought to be produced from the gene operonic with *tsuA* and contributes to the efficient assimilation of TS (Fig. 5B). Given that TsuB belongs to the TusA family of proteins, we predict that it is involved in guiding TS from TsuA to the first enzyme in assimilation, CysK. These observations mirror recent work in *E. coli* and, together, fortify our understanding of how bacteria access this inorganic metabolite to meet the nutritional sulfur requirement (25, 26, 60). Most importantly, however, is that these data validate the RNAseq by demonstrating that transcripts of genes with predicted sulfur metabolism function indeed contribute to meeting this nutritional requirement. Overall, this work provides compelling Datasets that highlight the *S. aureus* response to various states of sulfur supplementation and can be utilized to probe the nuances of sulfur metabolism to better understand this persistent threat to global health.

## ACKNOWLEDGEMENTS

We thank Dr. Jeffery Bose for supplying the pKK22 vector. Transposon mutants were acquired from the Network on Antimicrobial Resistance in *Staphylococcus aureus* (NARSA) for distribution by BEI Resources, NIAID, NIH, and the Nebraska Transposon Mutant Library (NTML) Screening Array NR-48501. We thank Drs. Scott Sherrill-Mix and Sean Crosson for assistance with the bioinformatics analysis. This work is funded by the National Institutes of Health R01 AI139074 and R21 AI142517.

## MATERIALS AND METHODS

### Strains and Primers

A complete list of strains and plasmids as well as primers used in this study can be found in Tables S3, S4, and S5 respectively. *S. aureus* JE2, a derivative of the community acquired USA300 LAC (68), was used as the WT strain. *Bursa aurealis* Tn inactivated strains were generated by transducing the Tn inactivated gene from the Nebraska Transposon Mutant Library (NTML) into JE2 (68–70). Tn insertions and chromosomal deletions were verified using PCR. Plasmids were confirmed via sequencing (Plasmidsaurus). Complementation studies were performed with the pKK22 vector (59). The pKK22-P*_tcyP_*-YFP fluorescence reporter plasmid was assembled using PCR-amplified YFP sequence from pAH462 (71, 72), 500 bp upstream of the *tcyP* gene (SAUSA300_RS02035), and the pKK22 vector via Gibson cloning.

### Media, growth conditions, and induction of sulfur starvation

Strains were routinely cultured overnight in tryptic soy broth (TSB; Remel) at 37°C, shaking at 225 rpm. CDM was prepared as previously described with slight modifications for the supplementation with the indicated sulfur sources (27, 28).

To quantify activity of the *tcyP* promoter activity using the YFP reporter, sulfur starvation was induced via the following method. Overnight TSB cultures of WT or *cymR*::Tn mutant cells harboring the pKK22-P*_tcyP_*-YFP plasmid were normalized to an OD_600_ of 1 and sub-cultured via a 1:100 dilution into CDM supplemented with 25 µM CSSC. Cells were incubated at 37°C with shaking (225 rpms) for 4 h. Upon reaching mid-exponential phase, the cells were collected via centrifugation and washed once in 1X phosphate buffered saline (PBS) followed by resuspension in fresh CDM supplemented with or without 25 µM CSSC and incubated for additional 2 h at 37°C with shaking (225 rpms). Aliquots (100 µL) were pipetted into triplicate wells of a black-walled 96 well plate (Corning) and monitored using a Biotek Synergy H1 Plate Reader (Agilent, Santa Clara, CA) for growth (OD_600_) and fluorescence (excitation at 488 nm and emission at 520 nm). Cells lacking the pKK22-P*_tcyP_*-YFP plasmid served as a control for background fluorescence.

For RNAseq experiments, independent biological duplicate cultures were prepared on distinct days. Sulfur starvation was induced by washing WT or *cymR*::Tn overnight cultures in PBS to a normalized optical density at OD_600_ of 1. Cells were then subcultured 1:100 into two 250 mL Erlenmeyer flasks containing 50 mL CDM supplemented with 25 µM CSSC. Flasks were incubated at 37°C, shaking at 225 rpm, for 4 h (Fig. S1). The duplicate flasks were then combined, centrifuged at 4°C, 4700 rpm for 10 min, washed with PBS, and resuspended in 100 mL fresh CDM lacking a sulfur source. The resulting resuspension was separated by pipetting 50 mL into two 250 mL Erlenmeyer flasks. One flask of WT and *cymR*::Tn was supplemented with 25 µM CSSC while the other remained sulfur deplete. Flasks were incubated at 37°C, shaking at 225 rpm, for 2 h prior to RNA isolation. To assess transcriptional changes between organic (Cys, CSSC, GSH, and GSSG) or inorganic (sTS) sulfur sources, WT cells were prepared identically as described above up to OD_600_ normalization. Cells were then subcultured 1:100 in CDM supplemented with either 50 µM Cys, 25 µM CSSC, 25 µM GSSG, 50 µM GSH, or 50 µM sTS, incubated at 37°C, 225 rpm, for 4 h prior to RNA isolation.

For growth curves, overnight cultures were washed in PBS and normalized to an OD_600_ equal to 1. Cells were diluted 1:100 into CDM supplemented with 25 µM CSSC, 50 µM sTS, 50 µM GSH, 750 µM GSH, 25 µM GSSG, 375 µM GSSG, or without a source of sulfur. Where described, a 30 mM hemin stock (dissolved in 1.4 M NH_4_OH) was diluted into the CDM at a final concentration of 10 µM. OD_600_ was monitored for 24 h in Epoch2 Biotek microplate spectrophotometer (Agilent) at 37°C, with continuous shaking. The growth curve for each strain represents the mean of three independent experiments, each conducted in biological triplicate.

### RNA extraction and sequencing

50 mL cultures from the respective growth conditions were centrifuged at 4°C for 10 min at 4700 rpm. RNA was isolated from the resulting pellet as previously described (22). The RNA was then treated with Turbo DNase following the manufacturer’s instructions (ThermoFisher, Waltham, MA). Total RNA was sent to Genewiz Inc. (South Plainfield, NJ) who conducted rRNA depletion with Illumina Ribo-Zero Plus (Illumina, Inc., San Diego, CA) and then generated libraries using stranded total RNA kit and sequenced using Illumina HiSeq (Illumina, Inc.) with 2x 150 bp paired-end read technology.

### Differential gene expression analysis and graphical representation

Gene expression was analyzed using Geneious Prime 2022.0.2 (Dotmatics, Boston, MA), with default parameters. Briefly, paired-end fastq files were trimmed using Bbduk plugin, and mapped and annotated against the reference genome *S. aureus* USA300_FRP3757 (NC_007793.1) using Geneious mapper. The expression levels were calculated within the software using DESeq2, which implements Wald test for initial hypothesis testing, followed by Benjamini-Hochberg for adjusted *P-*value calculations (73). Data was filtered using adjusted *P*-value < 0.05, and log_2_ FC (fold change) ≥ 1 or ≤ -1.

Functional gene categorization was performed evaluating cluster of orthologous groups (COG) categories with eggNOG-mapper v2 using protein sequences from *S. aureus* (NC_007793.1) (74, 75). Protein sequences that had no resulting COG assignment were denoted as “not classified”, and those not found in eggNOG-mapper were checked in EggNOG v6.0, using equivalent proteins in *S. aureus* NCTC8325, and characterized as “no homolog found” when no hits were detected. Enrichment analysis was performed using BioCyc, selecting Fisher-exact test with *P*-value < 0.05, and *S. aureus* USA300_FPR3757 as the reference (76).

### H_2_O_2_ peroxide killing assay

Using a modified protocol (77), overnight WT TSB cultures were washed in PBS and normalized to an OD_600_ of 1 and then subcultured 1:100 into two 250 mL Erlenmeyer flasks containing 50 mL CDM supplemented with either 50 µM GSH, 750 µM GSH, or 375 µM GSSG. Cultures were incubated at 37°C, shaking at 225 rpm, for 4 h. Cells were then transferred to a 50 mL falcon tube and spun at 4000 rpm and then resuspended in 50 mL PBS. 1 mL of the resuspension was utilized to obtain the OD_600_ while the remaining cells were spun at 4000 rpm. Cells were gently resuspended in residual PBS. One falcon tube for each growth condition was then resuspended in PBS to a final OD_600_ of 0.7. The other tube was resuspended to the same OD_600_ with PBS containing 1 M H_2_O_2_. Tubes were then placed at 37°C, static, for 20 min. Afterwards 3x 50 µL aliquots (i.e., 3 technical replicates) from each condition were resuspended in 950 µL PBS containing approximately 2,000-5,000 units mg^-1^ catalase (Sigma-Aldrich). Samples were then incubated at 37°C, static, for 5 min. Catalase inactivated samples were then serial diluted and incubated overnight at 37°C. Note that the average of these technical triplicates is considered as one biological replicate. Each assay was performed in biological triplicate.

### Figures and data availability

Figures were created using BioRender (BioRender.com). Sequenced RNA reads are available in NCBI (National Library of Medicine)-SRA (Sequence Read Archive) under BioProject PRJNA989516.

## Notes

### Competing Interest Statement

The authors have declared no competing interest.

### Summary of Updates

the supplemental files have been uploaded

