## Supplemental file for "Defining the transcriptional adaptation of *Staphylococcus aureus* to a range of nutritional sulfur supplementation"

**Supplemental Table 1. Genes upregulated in the *cymR* mutant in sulfur replete conditions**

| **Locus** | **Gene** | **Product** | **Log_2_ FC^a^** | **Adjusted *P*-value^b^** | **COG^c^** |
| --- | --- | --- | --- | --- | --- |
| SAUSA300_RS10985 |  | YeeE/YedE family protein | 3.435 | 5.131E-07 | S |
| SAUSA300_RS00910 |  | DUF4242 domain-containing protein | 3.430 | 1.335E-06 | S |
| SAUSA300_RS10980 |  | sulfurtransferase TusA family protein | 3.325 | 9.084E-07 | O |
| SAUSA300_RS00925 | *ssuC* | ABC transporter permease | 3.109 | 2.476E-06 | P |
| SAUSA300_RS00930 |  | acyl-CoA/acyl-ACP dehydrogenase | 3.059 | 5.169E-07 | I |
| SAUSA300_RS13915 | *isaA* | lytic transglycosylase IsaA | 2.807 | 1.255E-03 | M |
| SAUSA300_RS02035 | *tcyP* | l-cystine transporter | 2.699 | 1.428E-04 | U |
| SAUSA300_RS05050 |  | DoxX family protein | 2.369 | 6.815E-04 | S |
| SAUSA300_RS02045 |  | hypothetical protein | 2.290 | 2.041E-04 | A |
| SAUSA300_RS01055 | *gisA* | ABC transporter ATP-binding protein | 2.266 | 3.735E-03 | P |
| SAUSA300_RS02340 | *gmpC* | dipeptide ABC transporter glycylmethionine-binding lipoprotein | 2.194 | 6.966E-03 | P |
| SAUSA300_RS12610 | *sdpC* | CPBP family intramembrane glutamic endopeptidase SdpC | 2.104 | 2.280E-03 | S |
| SAUSA300_RS00915 | *ssuB* | ABC transporter ATP-binding protein | 2.053 | 1.266E-02 | P |
| SAUSA300_RS05345 |  | YkyA family protein | 2.048 | 1.382E-02 | L |
| SAUSA300_RS02635 | *cysK* | cysteine synthase A | 2.035 | 1.199E-02 | E |
| SAUSA300_RS09430 |  | hypothetical protein | 2.020 | 1.388E-02 | not classified |
| SAUSA300_RS02325 | *mccB* | bifunctional cystathionine gamma-lyase/homocysteine desulfhydrase | 1.988 | 2.250E-02 | E |
| SAUSA300_RS02335 | *gmpB* | methionine ABC transporter permease | 1.977 | 2.036E-02 | P |
| SAUSA300_RS09440 | *crcB2* | CrcB family protein | 1.976 | 9.513E-03 | D |
| SAUSA300_RS01875 |  | low temperature requirement protein A | 1.933 | 1.404E-02 | S |
| SAUSA300_RS13605 |  | ATP-binding cassette domain-containing protein | 1.906 | 1.654E-02 | V |
| SAUSA300_RS00920 | *ssuA* | ABC transporter substrate-binding protein | 1.904 | 2.249E-02 | P |
| SAUSA300_RS02330 | *gmpA* | methionine ABC transporter ATP-binding protein | 1.904 | 1.135E-02 | P |
| SAUSA300_RS12685 |  | alpha/beta hydrolase | 1.878 | 3.211E-02 | I |
| SAUSA300_RS01060 | *gisB* | ABC transporter permease | 1.834 | 3.649E-02 | EP |
| SAUSA300_RS13940 |  | DUF896 domain-containing protein | 1.825 | 4.623E-02 | S |
| SAUSA300_RS04485 |  | DUF3055 domain-containing protein | 1.822 | 1.364E-02 | S |
| SAUSA300_RS02855 |  | M20 family metallopeptidase | 1.785 | 2.250E-02 | E |
| SAUSA300_RS07460 |  | hypothetical protein | 1.725 | 2.352E-02 | no homolog found |
| SAUSA300_RS13025 | *tcyA* | transporter substrate-binding domain-containing protein | 1.693 | 2.663E-02 | ET |
| SAUSA300_RS13320 | *lnsB* | CPBP family lipoprotein N-acylation protein LnsB | 1.654 | 3.853E-02 | S |
| SAUSA300_RS10230 |  | hypothetical protein | 1.646 | 3.008E-02 | not classified |
| SAUSA300_RS05540 | *isdA* | LPXTG-anchored heme-scavenging protein IsdA | 1.646 | 3.382E-02 | M |
| SAUSA300_RS12705 |  | TetR/AcrR family transcriptional regulator | 1.620 | 4.532E-02 | K |
| SAUSA300_RS11450 |  | DUF2529 domain-containing protein | 1.618 | 2.554E-02 | S |
| SAUSA300_RS00605 | *sirA* | staphyloferrin B ABC transporter substrate-binding protein SirA | 1.608 | 2.782E-02 | P |
| SAUSA300_RS12440 |  | CHAP domain-containing protein | 1.588 | 3.896E-02 | S |
| SAUSA300_RS03340 | *tagA* | N-acetylglucosaminyldiphosphoundecaprenol N-acetyl-beta-D-mannosaminyltransferase TarA | 1.532 | 3.008E-02 | M |
| SAUSA300_RS07465 | *cmk* | (d)CMP kinase | 1.530 | 2.784E-02 | F |
| SAUSA300_RS04060 | *clpP* | ATP-dependent Clp endopeptidase proteolytic subunit ClpP | 1.525 | 4.532E-02 | OU |
| SAUSA300_RS00985 | *brnQ1* | branched-chain amino acid transport system II carrier protein | 1.503 | 4.983E-02 | E |
| SAUSA300_RS04205 | *lnsA* | lipoprotein N-acylation protein LnsA | 1.498 | 3.766E-02 | not classified |
| SAUSA300_RS07075 | *brnQ3* | branched-chain amino acid transport system II carrier protein | 1.469 | 4.532E-02 | E |
| SAUSA300_RS15260 |  | hypothetical protein | 1.454 | 4.532E-02 | no homolog found |
| SAUSA300_RS09900 |  | PTS transporter subunit IIC | 1.454 | 4.532E-02 | S |

^a^Expression ratio of *cymR*::Tn grown in medium containing CSSC relative to WT grown in the same medium.

^b^Adjusted *P*-value from DESeq2 output.

^c^COG assignments using eggNOG-mapper. A: RNA processing and modification; D: cell cycle control and mitosis; E: amino acid metabolism and transport; F: nucleotide metabolism and transport; I: lipid metabolism; K: transcription; L: replication and repair; M: cell wall/membrane/ envelope biogenesis; O: post-translational modification, protein turnover, chaperone function; P: inorganic ion transport and metabolism; S: function unknown; T: signal transduction; U: intracellular trafficking and secretion; V: defense mechanisms . Not classified: genes with an eggNOG-mapper homolog, but no associated COG. No homologs found: neither eggNOG-mapper nor Eggnog v6 could find a homolog for these proteins.

Genes highlighted in gray indicate overlap with Soutourina *et al*. (1).

**Supplemental Table 2. *S. aureus* sulfur starvation induces differential expression of genes encoding transcriptional regulators.**

| **Locus** | | | **Gene** | **description** | **Log_2_ FC^a^** | **Adjusted *P*-value^b^** | |
| --- | --- | --- | --- | --- | --- | --- | --- |
| **increased abundance** | | | | | | | |
| SAUSA300_RS00650 | *sbnI* | bifunctional transcriptional regulator/O-phospho-L-serine synthase SbnI | | | 2.787 | | 4.361E-11 |
| SAUSA300_RS04840 | *spxA* | transcriptional regulator SpxA | | | 2.710 | | 2.553E-09 |
| SAUSA300_RS07905 | *fur* | Fur family transcriptional regulator | | | 2.692 | | 9.202E-11 |
| SAUSA300_RS05125 |  | MarR family transcriptional regulator | | | 2.488 | | 6.488E-06 |
| SAUSA300_RS08020 | *argR* | transcriptional regulator ArgR | | | 2.348 | | 2.264E-07 |
| SAUSA300_RS12240 | *sarV* | HTH-type transcriptional regulator SarV | | | 2.319 | | 6.725E-06 |
| SAUSA300_RS03710 | *saeR* | response regulator transcription factor SaeR | | | 2.241 | | 2.563E-06 |
| SAUSA300_RS13930 |  | TetR/AcrR family transcriptional regulator | | | 2.009 | | 8.284E-06 |
| SAUSA300_RS03330 |  | metal-dependent transcriptional regulator | | | 1.998 | | 4.804E-06 |
| SAUSA300_RS12705 |  | TetR/AcrR family transcriptional regulator | | | 1.954 | | 5.729E-05 |
| SAUSA300_RS10810 |  | XRE family transcriptional regulator | | | 1.937 | | 5.507E-06 |
| SAUSA300_RS13640 |  | MarR family transcriptional regulator | | | 1.908 | | 1.274E-04 |
| SAUSA300_RS00590 | *sarS* | HTH-type transcriptional regulator SarS | | | 1.798 | | 1.302E-04 |
| SAUSA300_RS10675 |  | transcriptional activator RinB | | | 1.720 | | 1.793E-03 |
| SAUSA300_RS13480 | *scrA* | SaeRS system activator ScrA | | | 1.647 | | 1.974E-03 |
| SAUSA300_RS09835 |  | helix-turn-helix transcriptional regulator | | | 1.627 | | 3.238E-03 |
| SAUSA300_RS12900 |  | MerR family transcriptional regulator | | | 1.516 | | 3.018E-03 |
| SAUSA300_RS12510 |  | MurR/RpiR family transcriptional regulator | | | 1.506 | | 2.981E-04 |
| SAUSA300_RS01375 |  | GntR family transcriptional regulator | | | 1.481 | | 5.905E-04 |
| SAUSA300_RS08625 | *cymR* | Rrf2 family transcriptional regulator | | | 1.466 | | 1.744E-03 |
| SAUSA300_RS08905 | *nrdR* | transcriptional regulator NrdR | | | 1.432 | | 1.425E-03 |
| SAUSA300_RS06710 | *lexA* | transcriptional repressor LexA | | | 1.360 | | 1.701E-03 |
| SAUSA300_RS03605 | *mgrA* | HTH-type transcriptional regulator MgrA | | | 1.227 | | 9.413E-03 |
| SAUSA300_RS10805 |  | transcriptional regulator | | | 1.224 | | 1.972E-02 |
| SAUSA300_RS10800 |  | helix-turn-helix transcriptional regulator | | | 1.179 | | 2.229E-02 |
| SAUSA300_RS10060 | *perR* | peroxide-responsive transcriptional repressor PerR | | | 1.134 | | 2.713E-02 |
| SAUSA300_RS14275 |  | Crp/Fnr family transcriptional regulator | | | 1.131 | | 1.960E-02 |
| SAUSA300_RS12730 |  | MarR family transcriptional regulator | | | 1.032 | | 2.767E-02 |
| **decreased abundance** | | | | | | | |
| SAUSA300_RS06210 | *codY* | GTP-sensing pleiotropic transcriptional regulator CodY | | | -3.183 | | 3.373E-12 |
| SAUSA300_RS03665 |  | DeoR/GlpR family DNA-binding transcription regulator | | | -2.413 | | 5.295E-08 |
| SAUSA300_RS10935 | *agrB* | accessory gene regulator AgrB | | | -2.386 | | 4.644E-09 |
| SAUSA300_RS03250 | *sarA* | global transcriptional regulator SarA | | | -2.301 | | 1.061E-06 |
| SAUSA300_RS13560 |  | MerR family transcriptional regulator | | | -1.867 | | 4.348E-05 |

**^a^**Expression ratio of WT grown in sulfur deplete conditions relative to WT grown in sulfur replete conditions.

**^b^**Adjusted *P*-value from DESeq2 output.

**Supplemental Table 3.** **Strains used in this study.**

| Strain | Description | Reference |
| --- | --- | --- |
| wild type | USA300 LAC derivative JE2 | (2) |
| *cymR*::Tn | Nebraska Transposon Mutant Library (NTML) NE1293 *bursa* *aurealis* transposon (Tn) mutant, Erythromycin resistant (Erm^R^); SAUSA300_1583::Tn | (2) |
| *hrtA*::Tn | NTML NE1489 *bursa* *aurealis* Tn mutant, Erm^R^; SAUSA300_2306::Tn | (2) |
| SAUSA300_1998::Tn | NTML NE696 *bursa* *aurealis* Tn mutant, Erm^R^; *tsuA* | (2) |
| SAUSA300_0848::Tn | NTML NE966 *bursa* *aurealis* Tn mutant, Erm^R^; SAUSA300_RS04580 | (2) |
| *cymR*::Tn | *cymR* mutant backcrossed into JE2 | This study |
| *hrtA*::Tn | *hrtA* mutant backcrossed into JE2 | This study |
| *tsuA*::Tn | *tsuA* mutant backcrossed into JE2 | This study |
| SAUSA300_RS04580::Tn | Erm^r^ *Bursa* Tn insertion at JE2 chromosome position 927895 | This study |

**Supplemental Table 4.** **Plasmids used in this study.**

| Plasmid | Description | Reference |
| --- | --- | --- |
| pKK22 | Derivative of the naturally occurring *S. aureus* plasmid LAC-p01. Maintains stability in *S. aureus* without antibiotics. Linearized for Gibson assembly using primers PK85 & PK86. | (3) |
| pKK22-P*_tcyP_* -YFP | pKK22 encoding YFP under the control of the *tcyP* promoter. | This study |
| pKK22-*tsuAB* | pKK22 expressing *tsuA* (SAUSA300_RS10985) and *tsuB* (SAUSA300_RS10980) under native promoter expression. Insert generated with following primers: PK5 & PK8 | This study |
| pKK22-*tsuA* | pKK22 expressing *tsuA* (SAUSA300_RS10985) under native promoter expression. Generated insert with following primers: PK5 and PK10 | This study |
| pKK22-*tsuB* | pKK22 expressing *tsuB* (SAUSA300_RS10980) under native promoter expression. Insert generated as follows: PK5 and PK6 (promoter), PK7 and PK8 (*tsuB*); PK5 and PK8 used to sew promoter and *tsuB* fragments together before Gibson assembly. | This study |

**Supplemental Table 5.** **Primers utilized to generate strains in Table 1**

| Primer | Description | Sequence |
| --- | --- | --- |
| PK85 | pKK22 amplification/linearization for Gibson assembly | GCGGCCGCTAGCCTAGGAGC |
| PK86 | pKK22 amplification/linearization for Gibson assembly | ATCGCCTGTCACTTTGCTTGATATATGA |
| NE Martn-ermR* | Used to confirm Tn insertions on plus strand | CTCGATTCTATTAACAAGGG |
| NE Buster* | Used to confirm Tn insertions on minus strand | GCTTTTTCTAAATGTTTTTTAAGTAAATCAAGTAC |
| HL246 | Confirming *Bursa* Tn insertion in *cymR* | CTAATAACAAGATAACTTGACCAGAC |
| PK110 | Confirming *Bursa* Tn insertion in *hrtA* | AGAACTTAATGTCCCAGCC |
| HL267 | Confirming *Bursa* Tn insertion in *tsuA* | TCCCATACATATGCCACC |
| PK5 | Amplify *tsuAB* promoter and operon for Gibson assembly | CAAGCAAAGTGACAGGCGATTAGATGTTGTGATTCTAACTAC |
| PK6 | Amplify *tsuAB* promoter for Gibson assembly | CGTGTATCATAATCTTAACCTCTCATTTCC |
| PK7 | Amplify *tsuB* for Gibson assembly | GGTTAAGATTATGATACACGAATTAGGTAC |
| PK8 | Amplify *tsuB* for Gibson assembly | GCTCCTAGGCTAGCGGCCGCTTAAACTTTTTGAATTGTAATTGTC |
| PK10 | Amplify promoter and *tsuA* operon for Gibson assembly | GCTCCTAGGCTAGCGGCCGCCTATACTATTTGCGTTTGC |
| P*_tcyP_* forward | *tcyP* promoter region used for Gibson assembly | CAGGCGATGCGGCCGCGCCTAGAATTTTTTACAACGTGTTTGTTC |
| P*_tcyP_* reverse | *tcyP* promoter region used for Gibson assembly | TCATCGCTAGCACCTCCCTAGAACGTCACTCCTCAAATTTTTG |

*Reference: (2)

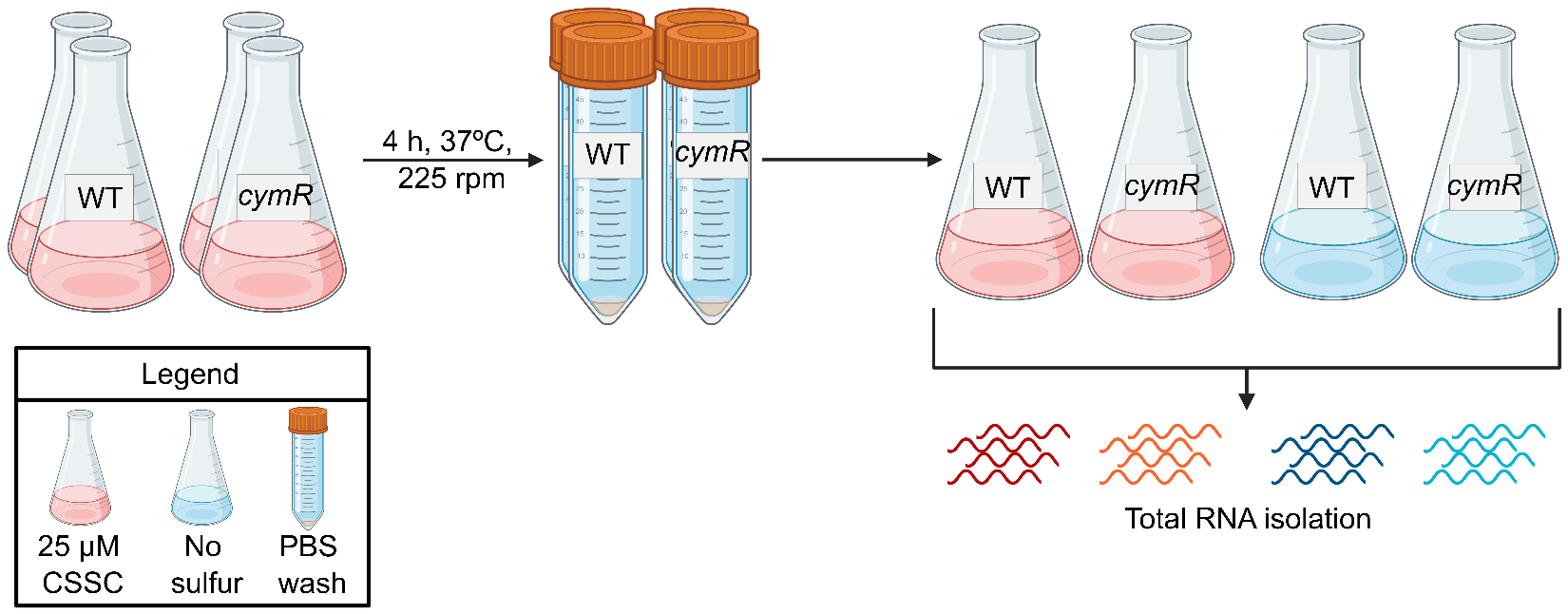

**Supplemental Figure 1. Experimental design employed to define the CymR-dependent and –independent responses to sulfur starvation in *S. aureus.*** WT or *cymR*::Tn were sub-cultured from normalized overnights into CDM supplemented with 25 µM CSSC and grown to mid-exponential phase (4 h). Cultures were pelleted, washed, and resuspended to the same cell density in medium with 25 µM CSSC or no viable sulfur source. Cells were grown for an additional 2 h prior to RNA isolation. Image generated with BioRender.

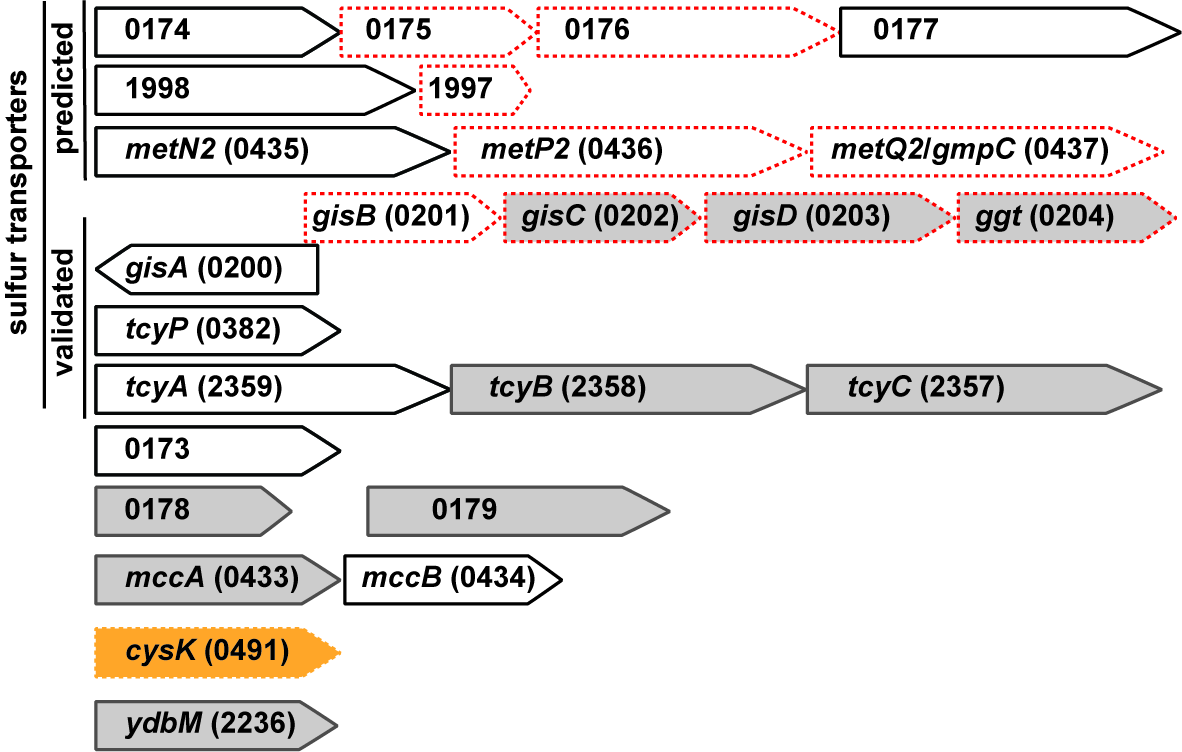

**Supplemental Figure 2. A refined *S. aureus* sulfur acquisition and metabolism gene repertoire based on comparative transcriptomics, operon organization, and genetic validation.** Transcripts of sulfur acquisition and metabolism genes that increase in abundance in response to WT sulfur starvation (Dataset 1), in the *cymR*::Tn mutant (Dataset 3), and were reported in Soutourina *et al*. (1) are presented. Genes that missed the statistical cutoffs (log fold change ≥2; *P*<0.05) in the sulfur replete *cymR*::Tn condition (Dataset 3) are gray. Genes that missed the statistical cutoff in the Soutourina study are denoted with a red dashed outline. *cysK* (orange) is the only gene that missed the statistical cutoff in the WT sulfur starvation condition (Dataset 1). Gene number refers to the SAUSA300_#### locus tag.

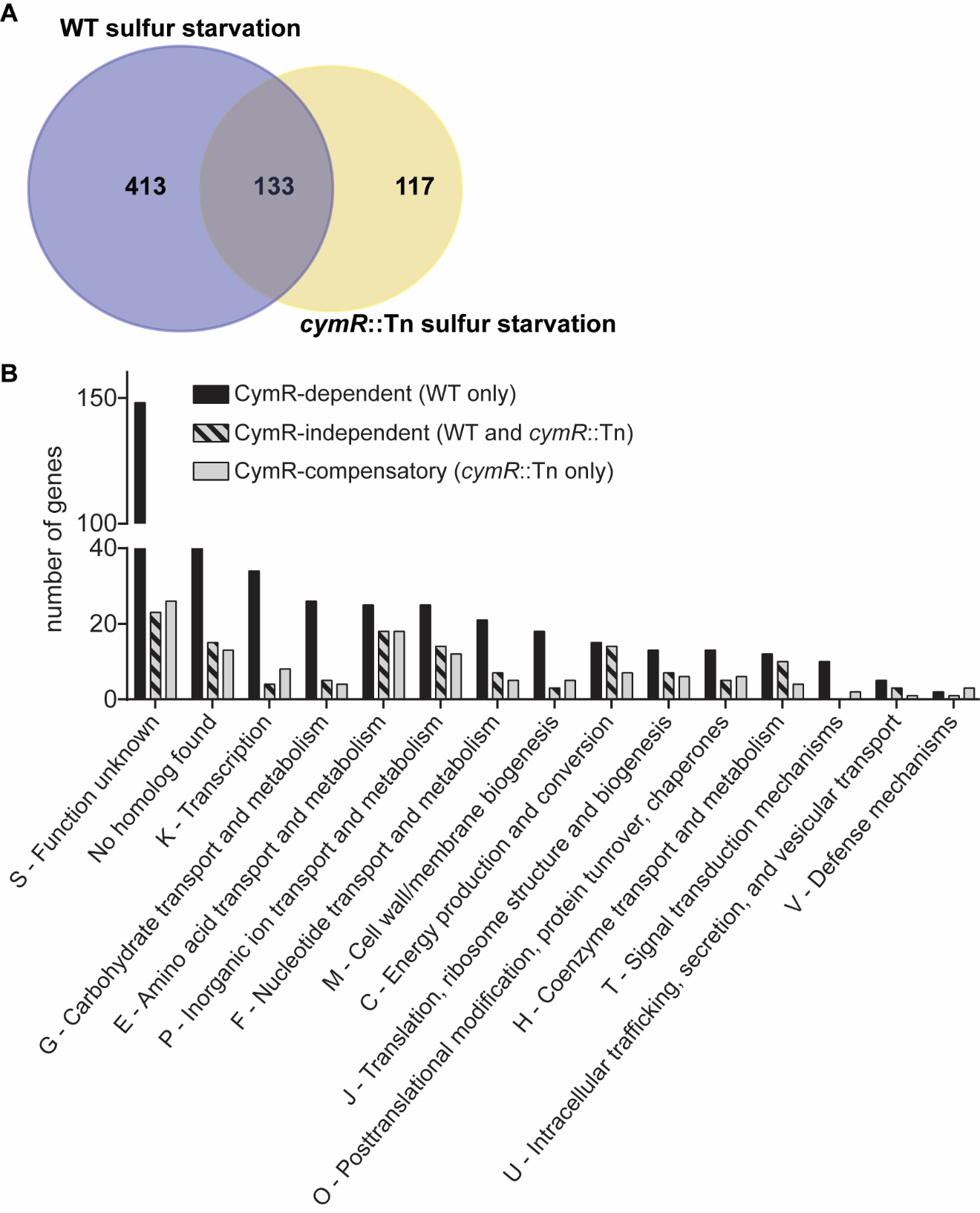

**Supplemental Figure 3. Comparison of CymR-dependent and -independent genes that respond to sulfur starvation. (A)** Number of genes that change in abundance in response to sulfur starvation in the presence (WT) or absence of CymR (*cymR*::Tn). WT sulfur starvation responsive loci from Table 1a (*n* = 546; blue) were compared against the starvation response when CymR is absent (Table 1c; *n* = 250; yellow). Genes that are shared between WT and the *cymR*::Tn mutant are considered to be CymR-independent (*n* = 133). CymR independent genes are those genes unique to the WT sulfur starvation condition (*n* = 413). Genes unique to the *cymR*::Tn condition (*n* = 117) are likely differentially expressed to compensate for the loss of CymR during sulfur starvation. **(B)** COGs assignment of genes in (A). eggNOG mapper was used to assign COGs (4-6). The number of COG assignments is greater than the number of genes in (A), as some genes were assigned multiple COGs.

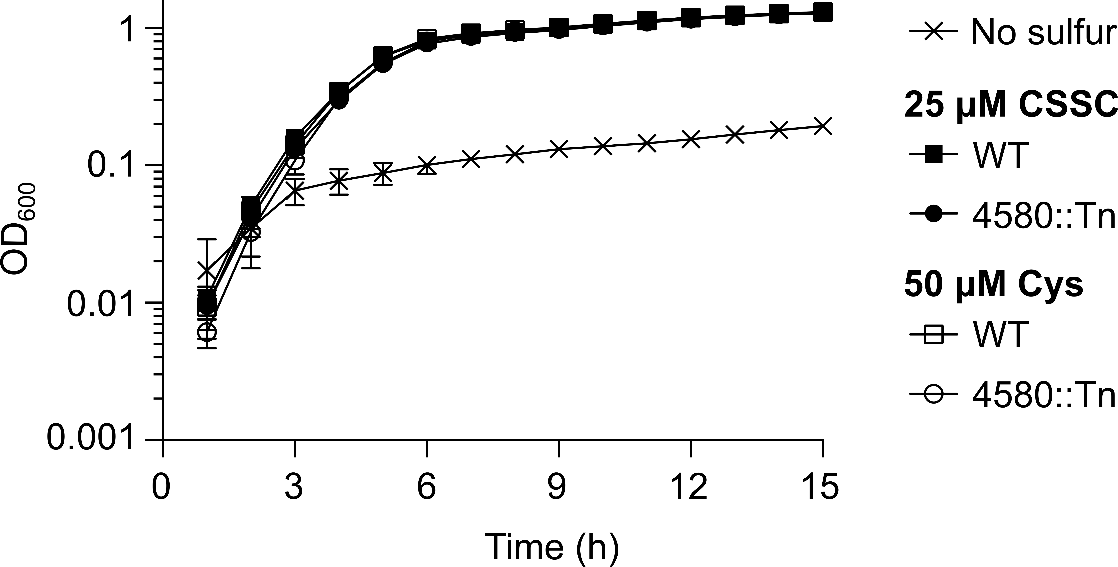

**Supplemental Figure 4. SAUSA300_RS04580 does not contribute to *S. aureus* proliferation on CSSC as a source of nutritional sulfur.** WT and SAUSA300_RS04580::Tn (4580::Tn) were cultured in chemically defined medium (CDM) supplemented with either 25 µM cystine (CSSC; closed symbols) or 50 µM cysteine (Cys; open symbols). Curves are the mean of three independent trials and the error bars depict ±1 standard error of the mean.
